# BioPhi: A platform for antibody design, humanization and humanness evaluation based on natural antibody repertoires and deep learning

**DOI:** 10.1101/2021.08.08.455394

**Authors:** David Prihoda, Jad Maamary, Andrew Waight, Veronica Juan, Laurence Fayadat-Dilman, Daniel Svozil, Danny A. Bitton

## Abstract

Despite recent advances in transgenic animal models and display technologies, humanization of mouse sequences remains the primary route for therapeutic antibody development. Traditionally, humanization is manual, laborious, and requires expert knowledge. Although automation efforts are advancing, existing methods are either demonstrated on a small scale or are entirely proprietary. To predict the immunogenicity risk, the human-likeness of sequences can be evaluated using existing humanness scores, but these lack diversity, granularity or interpretability. Meanwhile, immune repertoire sequencing has generated rich antibody libraries such as the Observed Antibody Space (OAS) that offer augmented diversity not yet exploited for antibody engineering. Here we present BioPhi, an open-source platform featuring novel methods for humanization (Sapiens) and humanness evaluation (OASis). Sapiens is a deep learning humanization method trained on the OAS database using language modeling. Based on an *in silico* humanization benchmark of 177 antibodies, Sapiens produced sequences at scale while achieving results comparable to that of human experts. OASis is a granular, interpretable and diverse humanness score based on 9-mer peptide search in the OAS. OASis separated human and non-human sequences with high accuracy, and correlated with clinical immunogenicity. Together, BioPhi offers an antibody design interface with automated methods that capture the richness of natural antibody repertoires to produce therapeutics with desired properties and accelerate antibody discovery campaigns.

BioPhi is accessible at https://biophi.dichlab.org and https://github.com/Merck/BioPhi.

## Introduction

Antibodies are versatile molecules that can bind diverse target antigens across the biological and chemical landscape. Antigen recognition is mediated by somatic hypermutation of the antibody variable region, where germline genes are combined and mutated to generate a diverse pool of mature sequences. Monoclonal antibodies (mAbs) represent the majority of protein-based therapeutics currently in the clinic, with clinical treatments in cancer [1], autoimmune disease [2], viral infection [3] etc. Commonly, mAbs are generated by the immunization of mouse or other model animal. However, sequences derived from rodent or other non-human sources are likely to elicit an immunogenic anti-drug antibody (ADA) response [4]. Therefore, the variable region of discovery mAbs must be humanized to mitigate undesirable clinical properties including safety risks and/or reduced efficiency. To do so, the hypervariable complementarity-determining regions (CDRs) and other essential murine framework residues are carefully incorporated into a human framework, producing a human-like sequence that preserves the binding properties of the original antibody. Although alternative antibody discovery approaches exist which avoid the need for humanization by employing transgenic mice with human B cell genes, this process is expensive and can still produce immunogenic sequences [5]. Additionally, human-like antibodies can also be developed inexpensively *in vitro* by high-throughput screening of large and diverse libraries using yeast or phage display technologies [6]. Nevertheless, antibodies produced in this manner have not undergone thymic selection, therefore they carry an immunogenicity risk and can further cross-react with other antigens. As a result of these caveats, mouse immunization and subsequent humanization of the murine sequences remains the primary path for antibody discovery.

Traditional humanization methods are based on germline sequences or natural sequence libraries of limited size. The canonical method of humanization is CDR grafting [7], by which the parental CDRs are inserted into a human germline sequence of choice. Additionally, positions important to the structural conformation of CDRs (known as “Vernier zones” [8]) are frequently back-mutated to the original parental residues. Although such residues can support the stability of the original CDR conformations, they can also reduce the effect of humanization. Expert knowledge is therefore needed to carefully balance this tradeoff, which limits the application to small numbers of sequences and is prohibitive to researchers who are lacking such expertise.

To evaluate the human-likeness of humanized sequences and identify immunogenicity risks, different “humanness scores” have been developed to date. First scores defined humanness based on sequence identity with a library of reference human sequences, averaged across all sequences in Z-score [9], or across the closest 20 sequences in T20 score [10]. A major pathway for the identification of foreign proteins is their processing into short peptides which are displayed on major histocompatibility complex (MHC) molecules and subsequently recognized by T-cell receptors. Guided by this principle, Human String Content (HSC) [11] derived a score from sequence identity of 9-mer peptides compared to sequences of human antibody germline genes. The HSC approach pioneered iterative humanization performed by maximizing the HSC score, later enabling joint optimization within the structural context [12]. Recently, the MG score approach [13] has enabled capturing higher-order relationships between pairs of sequence positions using a multivariate gaussian model, which was again applied as an optimization criterion for automated humanization. However, all of the abovementioned humanness scores have limited applicability for humanness evaluation due to the lack of granularity – a single score is provided for the entire sequence. Moreover, these scores are derived from small reference libraries, which can in turn impose unnecessary limitations on the diversity of the designed therapeutics.

The emergence of large-scale repertoires of natural antibodies provides a novel opportunity for exploiting antibody diversity to improve humanization and humanness evaluation methods. The Observed Antibody Space (OAS) database [14] has collected more than five hundred million human sequences from more than five hundred human subjects. Such repertoires not only inform our view of response and disease states, but also provide a diverse library of naïve and mature sequences that can be exploited for data-driven antibody engineering. The diversity and therapeutic applicability of OAS was demonstrated by its ability to recover sequences of therapeutic mAbs with high CDR sequence identity [15]. Furthermore, it was shown that developability properties of natural human antibodies are comparable with those of clinical mAbs [16]. In IgReconstruct [17], the OAS database was used to construct a back-translation method producing human-like DNA and evaluating humanness based on positional nucleotide frequency. Most recently, OAS was used in Hu-mAb [18] to train a random-forest-based humanness score used as an optimization criterion for iterative humanization.

The natural language processing field has demonstrated the ability of deep learning models to learn from enormous unlabeled bodies of text. Most recently, the Transformer architecture [19] has brought breakthroughs in question answering [20] or language translation [21]. Such progress is increasingly permeating the protein engineering space, where large-scale corpora of unlabeled data can likewise be exploited for real-world challenges. This has crystallized in AlphaFold [22], which demonstrated the applicability of large amounts of multiple sequence alignments (MSAs) and protein structures for protein folding prediction. Multiple methods were developed with the goal of encoding meaningful compact representations of protein sequences by pre-training on raw corpuses of amino acid sequences [23][24] or MSAs [25]. Such representations can be leveraged for transfer learning to specific problems with limited training sets [23]. In antibody discovery, AbLSTM [26] has demonstrated the ability of deep learning to distinguish between human and non-human antibody sequences. Deep learning was also able to select candidates with high affinity for iterative binding optimization using directed mutagenesis [27]. Recently, generative adversarial networks were trained on antibody repertoires to generate libraries of random antibody sequences with favorable developability profiles [28] or improved affinity [29].

Here we present BioPhi, a web platform for antibody design, humanization and humanness evaluation using natural antibody repertoires (Figure 1). Sapiens is a deep learning antibody humanization method trained on antibody repertoires of 266 human subjects from the OAS. Based on a test set of 25 humanized antibodies with known parental sequences and a novel test set of 152 humanized antibodies with putative parental sequences, we show that Sapiens produced humanization results comparable to those produced by expert methods. OASis (Observed Antibody Space identity search) is a novel antibody humanness score based on exact 9-mer peptide search in the OAS. OASis provides an interpretable and granular humanness report with adjustable stringency to allow evaluating sequences of high diversity. BioPhi is an open and extensible platform that offers improved design of humanized antibodies and will continue growing to facilitate faster discovery and development of antibody therapeutics.

**Figure 1:**
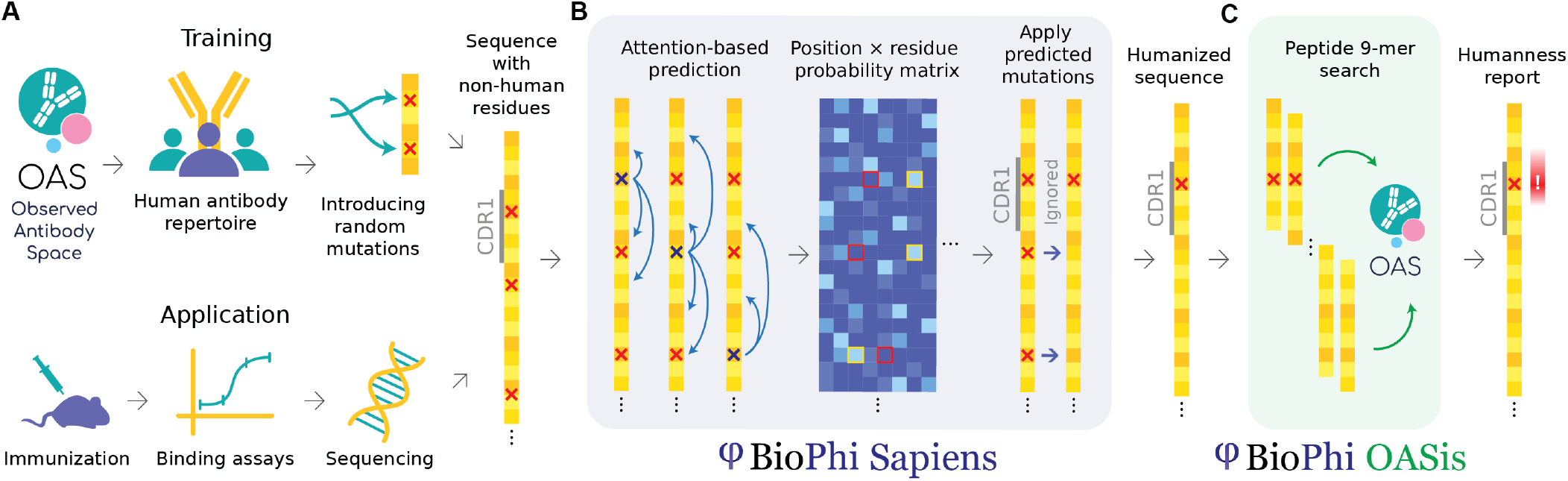
BioPhi integrated pipeline for bulk humanization (Sapiens) and humanness evaluation (OASis). (A) Sapiens is trained on human variable region antibody sequences from the OAS. Random positions in unaligned amino acid sequences are masked or mutated, Sapiens is trained to recognize and repair them. This simulates the real-world application where sequences with non-human residues originate from immunized mice, rabbit or other species. (B) Sapiens recognizes non-human residues using deep learning attention mechanisms and predicts the most probable human residues at each position given a particular input sequence, thereby humanizing it. (C) OASis evaluates humanness of an antibody sequence by chopping it into all overlapping 9-mer peptides and searching them against the OAS to estimate their prevalence across the human population

## Results

### Peptides from antibody repertoires capture diversity for antibody engineering

In antibody discovery and development, a large number of candidate variants need to be explored in order to find at least one lead candidate that satisfies the growing demands from developability [30], immunogenicity [4], or post-translational modification liability [31]. Therefore, a diverse antibody reference library needs to be collected and compactly represented in order to quickly and reliably determine which engineering mutations are viable and which do not conform to the acceptable human sequence space.

To capture the diversity of human antibodies, we used human antibody repertoires curated in the OAS database [14]. To visualize the sequence diversity, we sampled each OAS subject for complete variable region sequences from each V gene family. Next, we collected a germline reference of all 406 human heavy V genes and 206 light V genes from IMGT Gene-DB [32]. We embedded these into two-dimensional space using UMAP [33], placing sequences closer or further apart from each other based on sequence similarity. The dimensionality reduction revealed areas in sequence space that were densely covered by germline sequences as well as diverse areas that had no germline coverage (Figure 2A,B).

**Figure 2:**
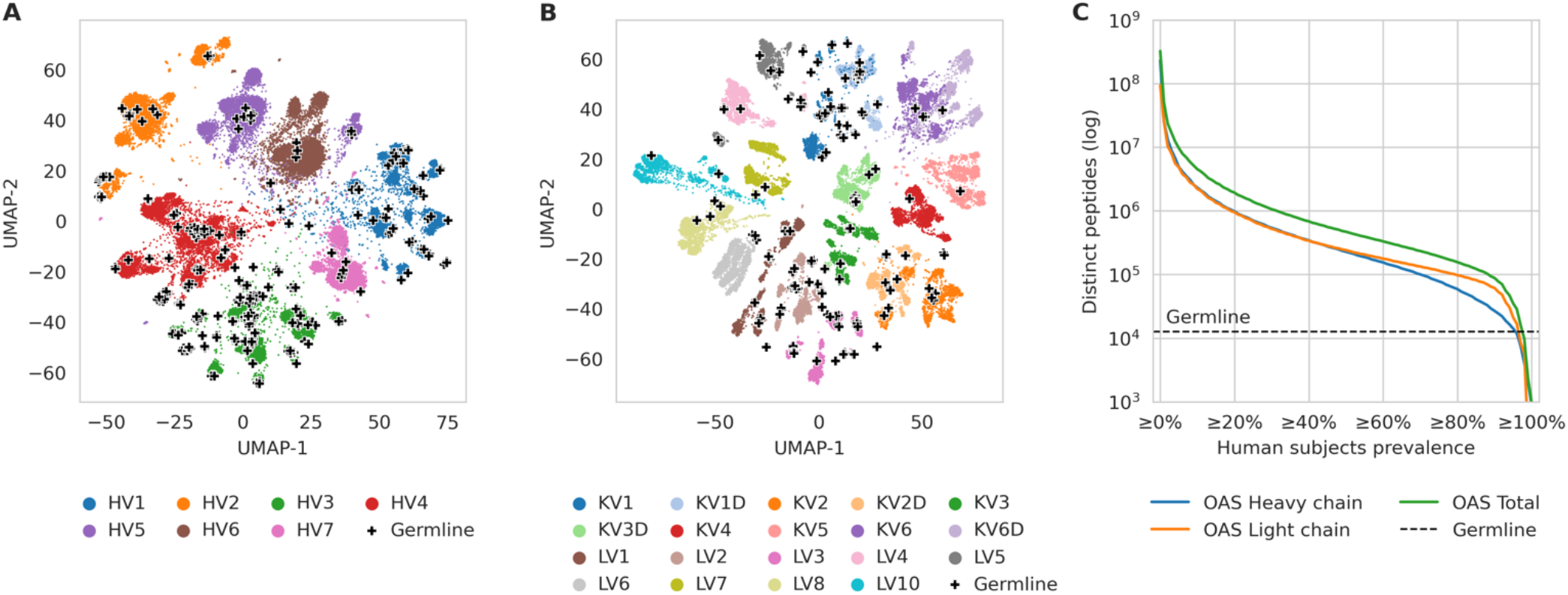
Diversity of human antibody germlines compared to human antibody repertoires from OAS. (A) and (B) Diversity of antibody variable region sequences illustrated using UMAP dimensionality reduction of approx. 40,000 variable heavy chain (A) and light chain (B) sequences randomly sampled from OAS. Each dot represents a complete variable region sequence (colored by germline family), each cross represents a germline V gene sequence. (B) Total number of distinct 9-mer peptides found in human germline compared to the number found in antibody repertoires across different fractions of human population, calculated from repertoires of 231 human subjects in OAS. For each fraction of subjects on the X axis, the Y axis shows the number of peptides that appear at least in the given fraction of subjects (e.g. there were 1.9×10^6^ distinct peptides that appeared in at least 20% of subjects).

Inspired by HSC score [11], which relies on 9-mer peptides from antibody germline, we relied on 9-mer peptides from antibody repertoires to capture a compressed representation of the wider antibody sequence diversity. We extracted all overlapping 9-mer peptides from variable heavy and light chains found in OAS repertoires that were linked to a single human subject (donor of the sample) and contained at least 10,000 complete sequences. To reduce sequencing errors and manage the peptide database size, we removed heavy chain peptides found in only one human subject. This yielded the *OASis* database comprising of 139,000,294 distinct peptides found across 118,713,869 sequences from 231 subjects from 26 studies in OAS. For each distinct peptide, we stored the list of corresponding subjects. These records enabled calculating the *prevalence* of any 9-mer peptide across the human population, which we defined as the percentage of subjects in OAS that contained an exact match of the peptide in at least one of the sequences in their antibody repertoire.

Next, we used the OASis database to estimate the total number of distinct 9-mer peptides shared across different fractions of the human population, in other words the public repertoire diversity. We compared our public repertoire diversity estimate with the number of distinct 9-mer peptides extracted from all human germline VDJ genes from IMGT Gene-DB. Even when considering only ubiquitous peptides found in more than 80% of human subjects in OAS, repertoires provided a 12-fold larger sequence space than germline in terms of the number of unique peptide “building blocks” (Figure 2C). Furthermore, this is a lower estimate since current sequencing depth is limited compared to estimated repertoire sizes of 10^11^ or more unique sequences [34].

In summary, the OASis peptide database offers diversity and granularity that can guide antibody engineering efforts with respect to humanization and immunogenicity decisions.

### OASis provides a granular and interpretable humanness evaluation

In order to provide utility for antibody engineering, the output of a humanness evaluation needs to be designed with both granularity and interpretability in mind. Granularity can be achieved by identifying how different residues or stretches of the sequence contribute to the overall humanness, as opposed to providing only one score for the whole sequence. Interpretability can be achieved by explaining the humanness score of a particular sequence comprehensively to the user, for example by near reference sequence matches, as opposed to black-box predictions.

To tackle these challenges, we developed OASis, a humanness evaluation method that reports human prevalence of individual peptides together with a single aggregated score for an antibody as a whole. First, OASis evaluates the humanness of each overlapping 9-mer peptide in an antibody sequence in terms of human prevalence (Figure 3A). Such evaluation provides a granular and interpretable humanness report that highlights regions that present the largest risk.

**Figure 3:**
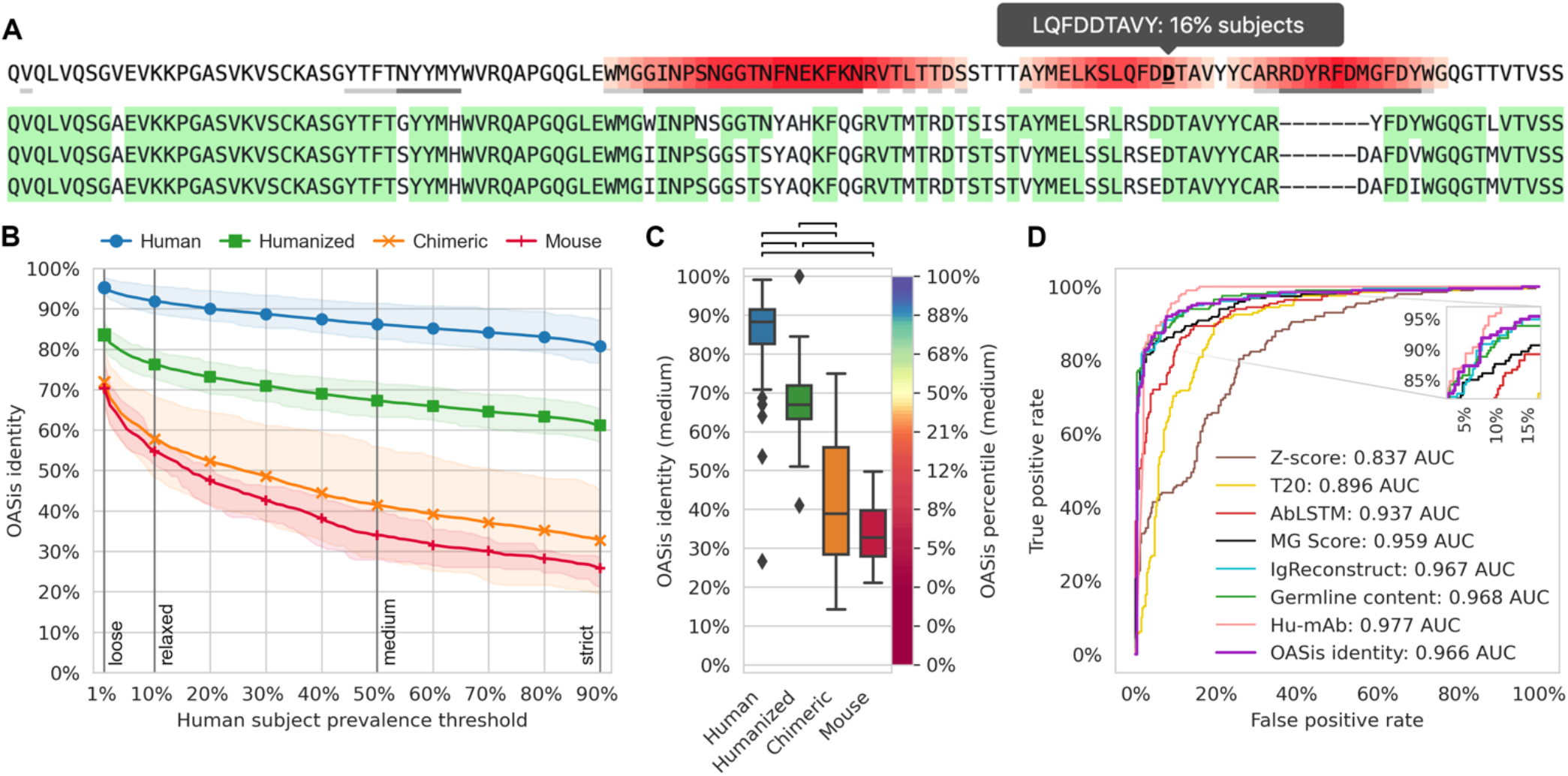
OASis provides a granular and interpretable humanness score that is able to separate therapeutic antibodies of different origin. (A) Example OASis peptide prevalence report of Pembrolizumab heavy chain sequence illustrating the granularity of OASis humanness evaluation. Red gradient corresponds to number of overlapping 9-mer peptides at given position that are identified as non-human based on user-specified prevalence threshold. Grey tooltip illustrates how prevalence is reported interactively on mouse hover. Three nearest germline sequences are shown below, green background marks sequence identity. (B) Ability of OASis identity to separate between human, humanized, chimeric or murine antibodies across different prevalence thresholds. The X axis shows the fraction of human OAS subjects needed to contain a peptide in order for it to be considered human, spanning from a loose prevalence of at least 1% subjects to a strict prevalence of at least 90% subjects. The Y axis shows the fraction of peptides in an antibody that are considered human at the given threshold (OASis identity). Lines show the average of each group, highlighted regions span between 25% and 75% quantiles. (C) Ability of OASis identity to separate between human, humanized, chimeric and murine antibodies at the *medium* threshold (50% prevalence). Left Y axis shows OASis identity, right Y axis shows OASis percentile (percentile of OASis identity among therapeutic antibodies). Black brackets show significant differences (p < 1e-8 based on two-sided Mann-Whitney U test). (D) Evaluation of OASis medium identity and other humanness metrics’ ability to distinguish between human (positive) and nonhuman (negative) therapeutic antibodies visualized using a ROC curve.

In order to evaluate and compare humanness at whole antibody level, we define the *OASis identity* score. OASis identity is calculated for a single antibody as the fraction of its peptides that pass a user-defined prevalence threshold. The prevalence threshold determines what fraction of the human population should contain a given peptide in order for it to be considered human. We made the threshold adjustable since we anticipate that future immunogenicity research will improve our understanding of how prevalent a given peptide needs to be in order to be considered safe. In the BioPhi platform, the threshold is fully customizable, but we also provide four pre-defined thresholds that capture different stringency levels: *loose* (≥ 1% subjects), *relaxed* (≥ 10% subjects), *medium* (≥ 50% subjects) and *strict* (≥ 90% subjects). For example, at the loose threshold, each peptide is identified as human if it is found in at least 1% of subjects. And correspondingly, the OASis loose identity score of an antibody is calculated as the fraction of its peptides that are found in at least 1% of subjects.

To facilitate easier interpretation of a particular OASis identity score, we further define the *OASis percentile* which converts the identity score to the 0-100% range based on 544 therapeutic mAbs from IMGT mAb DB [35]. Then, the 0% OASis percentile score corresponds to the least human and the 100% OASis percentile score corresponds to the most human antibody in the clinic. In BioPhi, humanness is reported using OASis as well as using traditional methods based on positional residue frequency and germline sequence identity (Supplementary Figure 1,2).

A humanness score should be able to distinguish between human antibodies and antibodies from other species. In particular, it should enable doing so for therapeutic antibodies since those are the primary subjects of humanness analysis. Therefore, we evaluated the ability of OASis identity to separate between antibody therapeutics of different origin extracted from IMGT mAb DB. For each group of 198 human, 229 humanized, 63 chimeric and 13 murine sequences, we calculated the average OASis identity across all 1%–90% prevalence thresholds (Figure 3B). These curves visualized humanness of each group of sequences across all prevalence thresholds, enabling humanness evaluation with respect to any definition of what prevalence is considered human. As expected, with increasing stringency, the curves demonstrated a decreasing humanness trend. This was especially visible in murine sequences where 70% peptides were considered human when prevalence in 1% subjects was required, while only 26% peptides were considered human when prevalence in 90% subjects was required.

We further visualized OASis identity at 50% prevalence which confirmed significant differences in the score distribution between different species (p < 1e-8, Figure 3C). Interestingly, two outliers with low humanness were identified in the human group, at 27% and 54% OASis identity score. The former was Elipovimab that targets HIV, the latter was Navivumab that targets Influenza A. Both mAbs scored low due to their long CDR3 sequences and heavily mutated frameworks that gave rise to many non-human 9-mer peptides identified across the variable region.

Next, we defined the humanness prediction problem as a classification task, where the positive class contained human sequences and negative class contained sequences that were humanized or from other species. For different humanness evaluation methods, there was little consensus on what score needed to be achieved to considered a sequence human. Therefore, we did not measure performance at a single score cutoff but used the standard receiver operating characteristic (ROC) curve approach that shows how the true positive rate (TPR) changes with the false positive rate (FPR) for all different score cutoffs (Figure 3D, Supplementary Figure 3). In this context, the TPR was the fraction of human sequences correctly predicted as human and FPR was the fraction of non-human sequences incorrectly predicted as human. OASis medium identity outperformed (0.966 AUC) the results of traditional humanness scores based on homology – Z-score (0.837 AUC) and T20 score (0.896 AUC) as well as two novel humanness scores – AbLSTM (0.937 AUC) and MG score (0.959 AUC). OASis performed comparably to IgReconstruct (0.967 AUC), a recent method based on back-translation and positional nucleotide frequency. Interestingly, OASis performance was also comparable to Germline content (0.968 AUC), a baseline score we implemented based on percent sequence identity with nearest human germline. All methods were outperformed by Hu-mAb (0.977 AUC), a recent random forest method trained on the particular task of classifying human and non-human sequences. However, IgReconstruct score is not granular and Hu-mAb score is neither granular nor interpretable. A comparative analysis is provided in Table 1. All scores and the 553 sequences used to produce them are provided in Supplementary Table 1.

**Table 1:** Evaluation of antibody humanness scores. A granular score reports how different residues or stretches of the sequence contribute to the overall humanness. An interpretable score explains the humanness of a particular sequence comprehensively to the user, for example by near sequence matches or prevalence statistics. Diversity is reported as the size of the reference sequence library (in orders of magnitude). Separately for each column, values above 75% percentile are marked green, values below 25% percentile are marked red.

| Method | Granularity | Interpretability | Diversity (seqs) | Humanness classification |  | Immunogenicity |  |
| --- | --- | --- | --- | --- | --- | --- | --- |
|  |  |  |  | Accuracy (%) | ROC AUC | R | R2 |
| Z-score | ✗ no | ✓ yes | 10<sup>3</sup> | 76.5 | 83.7 | -0.45 | 0.20 |
| T20 | ✗ no | ✓ yes | 10<sup>4</sup> | 83.9 | 89.6 | -0.41 | 0.16 |
| AbLSTM | ✗ no | ✗ no | 10<sup>4</sup> | 87.8 | 93.7 | -0.47 | 0.22 |
| MG Score | ✗ no | ✗ no | 10<sup>3</sup> | 91.8 | 95.9 | -0.46 | 0.21 |
| IgReconstruct | ✗ no | ✓ yes | 10<sup>8</sup> | 92.6 | 96.7 | -0.45 | 0.20 |
| Germline content | ✓ yes | ✓ yes | 10<sup>2</sup> | 92.8 | 96.8 | -0.50 | 0.25 |
| Hu-mAb | ✗ no | ✗ no | 10<sup>7</sup> | 93.3 | 97.7 | -0.58 | 0.34 |
| OASis identity (loose) | ✓ yes | ✓ yes | 10<sup>8</sup> | 91.9 | 96.4 | -0.48 | 0.23 |
| OASis identity (relaxed) | ✓ yes | ✓ yes | 10<sup>8</sup> | 94.4 | 97.2 | -0.51 | 0.26 |
| OASis identity (medium) | ✓ yes | ✓ yes | 10<sup>8</sup> | 92.6 | 96.6 | -0.53 | 0.28 |
| OASis identity (strict) | ✓ yes | ✓ yes | 10<sup>8</sup> | 91.9 | 95.6 | -0.53 | 0.28 |

Ultimately, the goal of humanness evaluation is to capture and reduce the immunogenicity risk. Thus, we evaluated the ability of OASis and other humanness metrics to predict reported anti-drug antibody response of 217 therapeutics curated in [18]. We measured performance using Pearson correlation coefficient (R) and explained variance (R^2^). OASis medium identity has outperformed (R = −0.53, R^2^ = 0.28, Supplementary Figure 4A) all results except those of Hu-mAb (R = −0.58, R^2^ = 0.34, Supplementary Figure 4B). Although germline content was also predictive of immunogenicity (R= −0.50, R^2^ = 0.25), the correlation was comparable to repertoire-based methods, supporting the assumption that the augmented diversity provided by natural antibody repertoires does not compromise the immunogenicity profile.

**Figure 4:**
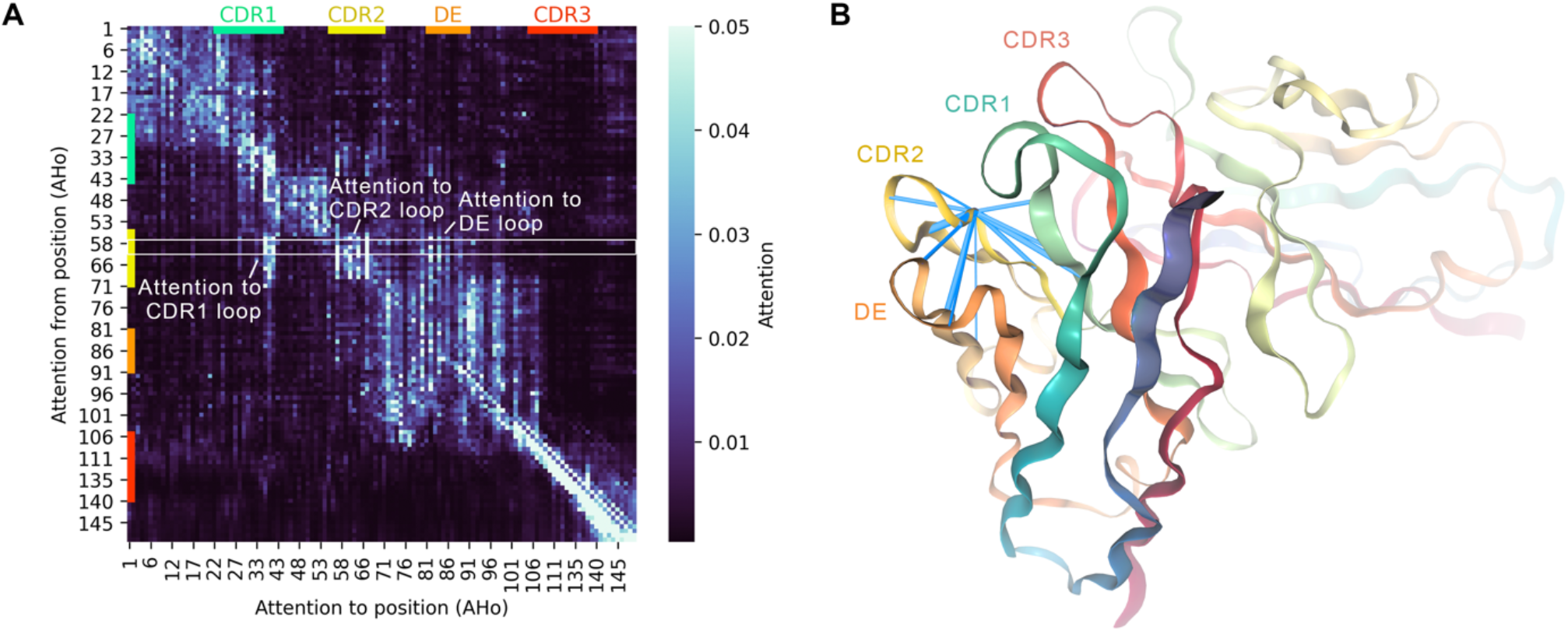
Attention within the Sapiens neural network captures long-range dependencies between antibody loops. (A) Attention weights of Sapiens when predicting Pembrolizumab heavy chain. Heatmap shows average of all attention heads in layer 2/4 of the Transformer encoder. The matrix defines the contribution of all positions (columns) when predicting the residue at a given position (row) in the sequence. White rectangle highlights attention from AHo position 59 visualized in (B). (B) Visualization of attention from Asparagine on AHo position 59 within CDR2 loop of Pembrolizumab 3D structure (PDB 5B8C). Blue beams visualize all attention connections with weight over 0.02, diameter proportional to weight.

Since immunogenicity is largely supported by the display of peptides on MHC-II receptors, we additionally evaluated a novel metric equal to the number of peptides that were predicted to bind MHC-II and that were not of human origin based on OASis. However, this has decreased the performance (R = 0.44, R^2^ = 0.01, Supplementary Figure 4C), therefore we have not pursued this further. All scores and the 217 sequences used to produce the data are provided in Supplementary Table 2.

**Table 2:**
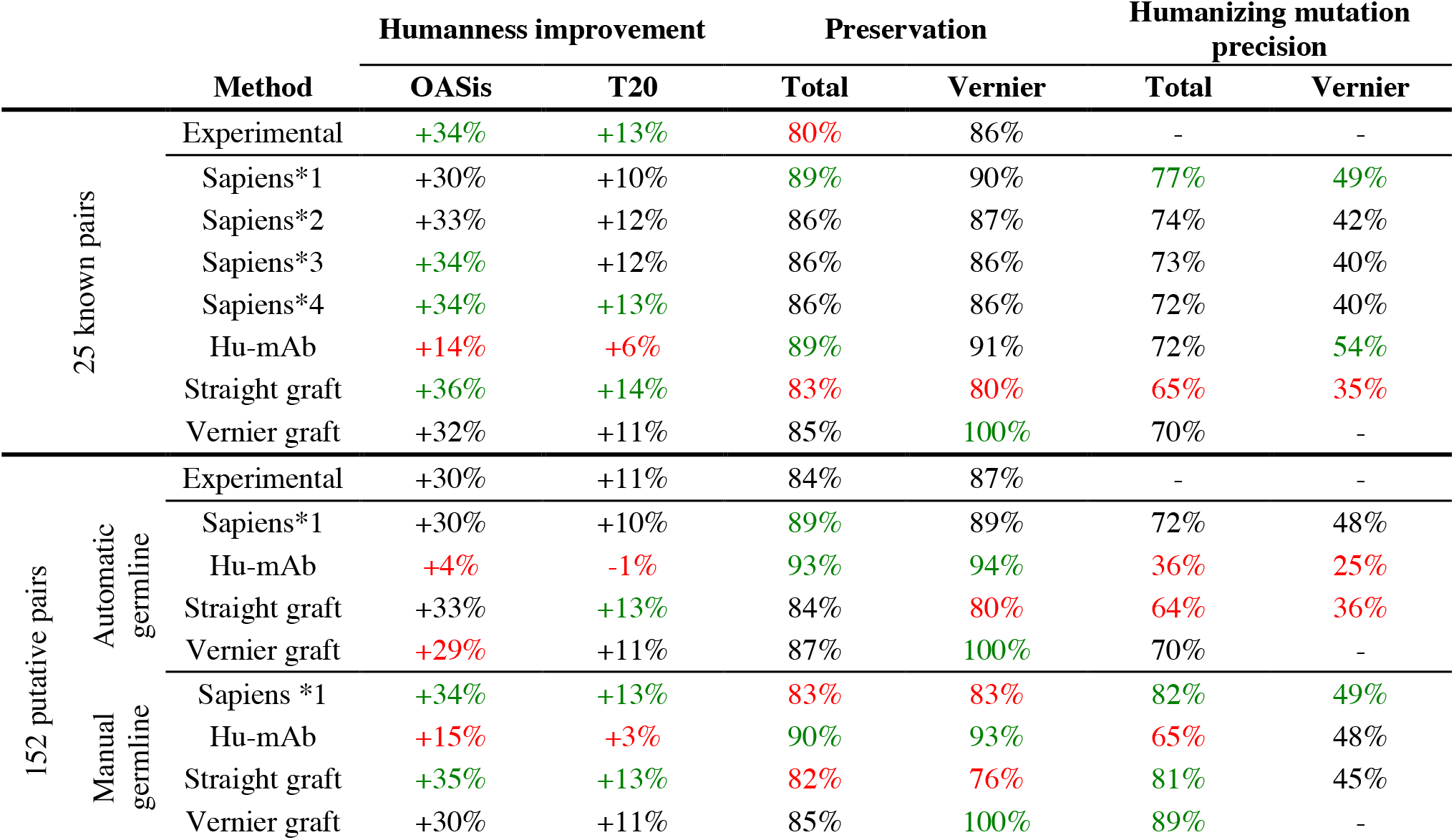
Evaluation of humanization methods. Separately for each column, values above 75% percentile are marked green, values below 25% percentile are marked red. Humanness change was computed as average absolute difference of OASis medium identity (or T20 score) of the humanized sequence and the parent sequence. Therefore, an increase of +34% refers to the absolute change in the humanness score (e.g. from 40% to 74%), not a relative change. Parental preservation was calculated as sequence identity of the parental and humanized sequence under Kabat numbering, in full sequence or Vernier regions only. Humanizing mutation precision was calculated as number of mutations made both in predicted sequence and in experimentally humanized sequence, divided by total number of mutations made in the predicted sequence.

In summary, OASis was able to estimate the humanness of a sequence with high accuracy and detect the risk of immunogenicity. When granularity and interpretability is required, OASis outperforms all existing humanness scores.

### Sapiens learns to represent antibody sequences using language modeling

Although peptides capture the nature of human antibodies as it relates to immunogenicity, they do not capture the long-range dependencies between positions, while these are important for structural stability and other general viability properties of a sequence. Therefore, we devised a separate humanization mechanism that can capture humanness including long-range dependencies. Moreover, such separation of the humanization method from the humanness score ensures independent humanness evaluation, in contrast to optimizing and evaluating using the same score as employed by previous approaches [11][13][18].

We developed Sapiens, a deep neural network based on the Transformer encoder architecture [19]. The Sapiens training procedure was based on masked language modeling [20], where the input “sentences” were amino acid sequences of antibody variable regions and the “word” tokens were individual amino acid residues. During training, some of the amino acids in the input sequence were randomly replaced with a mask token or mutated to a random amino acid. The model was trained to recognize these replacements and predict the original amino acids. The positions of the perturbed residues were not revealed to the model, therefore it needed to learn to recognize and repair any unexpected residues based on the context. Consequently, the model was trained only on human antibody sequences and no additional labelling data was needed. We trained a separate heavy chain and light chain model on a subset of 20 million heavy chain sequences and 19 million light chain sequences from the OAS (see Methods).

Compared to convolutional or recurrent neural networks, the Transformer neural network used by Sapiens is built exclusively on *attention* mechanisms. Analogous to language modeling where attention weights reveal dependencies between words in a sentence, in Sapiens they reveal dependencies between residues in an antibody sequence. In a traditional machine learning context where residues on different positions in a sequence are considered input features, attention can be thought of as feature importance, with two important distinctions. First, unlike feature importance, attention is not fixed but changes according to the input sequence. Second, attention can be calculated for a single position (importance of position A) as well as for a pair of positions (importance of position A in predicting position B). Attention is defined for a given input sequence using an *attention matrix*, which contains the importance of each position (column) when predicting the residue at each position (row).

By inspecting the attention patterns, we evaluated the ability of Sapiens to capture long-range dependencies. To calculate an average attention matrix, we selected heavy chain sequences from IMGT mAb DB that had exactly 120 residues and were composed of the same positions under AHo [36] numbering. Although the average attention matrix did not reveal many long-range residue contacts, some patterns were clearly visible (Figure 4A, Supplementary Figure 5). First, we observed that positions in framework regions 1-4 were directing their attention mostly to their respective region compared to other regions (3.8, 3.7, 4.4 and 15.0 times higher attention on average in framework 1-4 respectively). Second, we observed that positions in CDR2 loop were directing their attention towards the CDR1 and CDR2 loops as well as the DE loop compared to framework regions (2.4, 8.0 and 3.2 times higher on average for CDR1, CDR2 and DE respectively). To illustrate the spatial proximity of these loop positions, we calculated attention weights from Asparagine on AHo position 59 of Pembrolizumab heavy chain and visualized these in the crystal structure (Figure 4B).

**Figure 5:**
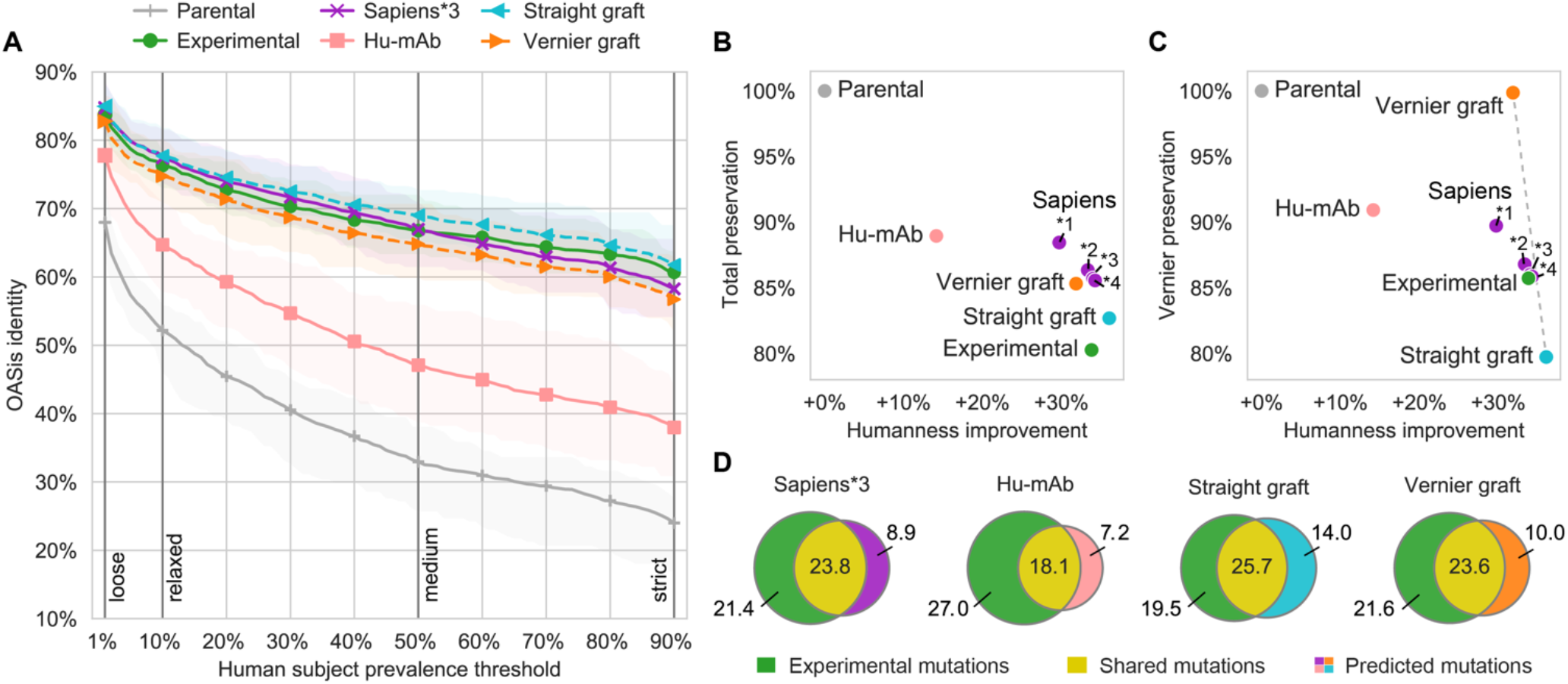
Sapiens achieved a balanced humanness-preservation tradeoff on 25 antibodies with known parental sequences. (A) Humanness of parental and humanized sequences evaluated using OASis identity curves. Each curve shows average across the 25 sequences, highlighted area spans between 25% and 75% quantiles. (B) Humanness-preservation tradeoff. X axis shows humanness improvement – average difference in OASis medium identity of the humanized and parental sequence. Y axis shows total parental sequence preservation (average percent sequence identity of humanized and parental sequence). Sapiens*1,*2,*3 and *4 refer to 1, 2, 3 and 4 humanization iterations respectively. (C) Humanness-preservation tradeoff in Vernier zones. X axis same as in (B), Y axis shows parental sequence preservation in Vernier zones. Dashed gray line shows axis between two extremes: Straight CDR graft (all Vernier residues humanized, more human but less preserved) and Vernier CDR graft (all Vernier residues back-mutated, less human but more preserved). (D) Average overlap of humanizing mutations made in predicted sequence and in experimentally validated sequence.

Since we observed increased attention between antibody loops that are not neighboring in the sequence but in the three-dimensional space, we thus concluded that Sapiens demonstrated the ability to capture long-range dependencies in antibody sequences.

### Sapiens achieves a balance of humanness and parental sequence preservation

The challenge of humanization is improving humanness while preserving as much of the parental sequence as possible in order to preserve binding and affinity. Ultimately, the only way to evaluate reduced immunogenicity risks and preserved binding affinity and functional activity is through *in vitro* or even *in vivo* assays. Nevertheless, *in silico* evaluation of humanization methods is necessary at least as a primary filter that enables comparing and selecting between multiple humanized candidates.

In this study, we define the *humanness-preservation tradeoff* as a way to evaluate the mutually competing goals of the increase in humanness and the preservation of the parental sequence. The first component, humanness increase, was measured as the absolute difference in OASis identity score of the humanized sequence compared to the parental sequence. The second component, parental sequence preservation, was measured as percentage sequence identity between the parental and humanized sequence overall (*Total preservation*) and in Vernier zones (*Vernier preservation*).

Since many humanized antibodies have been developed and tested, these can be used as standards for comparison. Primarily, their *parental* sequences (original sequences from mouse or other model organism) are needed in order to be able to produce and evaluate alternative humanized sequences. To that end, 7 pairs of parental-humanized sequences were first curated in [13] and recently expanded to 25 pairs in [18]. Thus, we used these 25 experimentally validated pairs as our first humanization benchmark. Analogous to existing approaches, we measured performance in terms of overlap between the humanizing mutations made in the predicted and the experimentally validated sequence. Since a sequence can be successfully humanized in various ways, even across different germlines, we could not consider the experimentally validated sequence as a single ground truth. Nevertheless, by highlighting the level of agreement between the prediction and a human expert, we provided an additional layer of confidence in our humanization method.

To define naïve baselines of automated humanization within the humanness-preservation context, we implemented two humanization methods based on CDR grafting. A *Straight* CDR graft was created by inserting the Kabat CDR regions into nearest human germline frameworks. Hence, this baseline prioritized humanness over parental sequence preservation. A *Vernier* CDR graft was created from the Straight CDR graft by additionally back-mutating all Vernier zone positions to the parental residues. Hence, this baseline prioritized parental sequence preservation over humanness improvement. An illustration is provided in Supplementary Figure 6.

**Figure 6:**
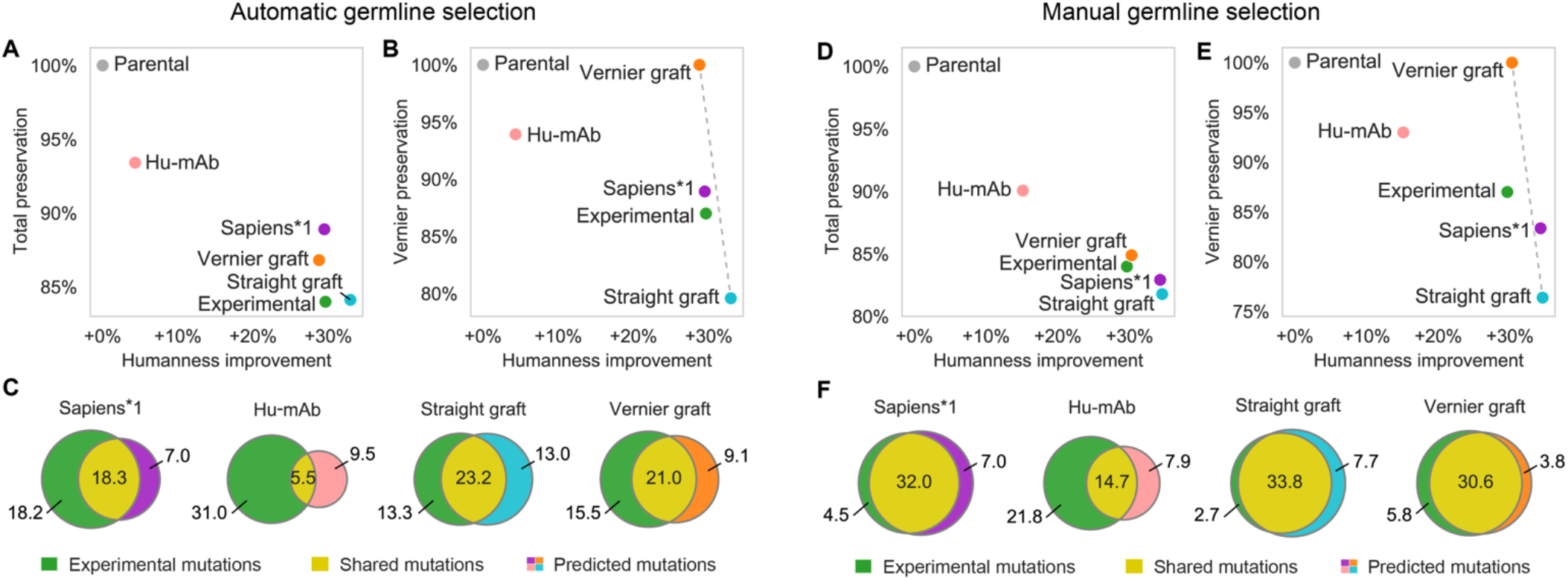
Evaluation of humanization methods on a large scale using 152 humanized therapeutic antibodies with putative parental sequences. In automatic germline selection (A-C), target germlines were chosen by the humanization method. In manual germline selection (D-F), target germline was set based on the germline of the humanized therapeutic sequence. (A) and (D) Humanness-preservation tradeoff. X axis shows humanness improvement – average difference in OASis medium identity of the humanized and parental sequence. Y axis shows total parental sequence preservation – average percent sequence identity of humanized and parental sequence. (B) and (E) Humanness-preservation tradeoff in Vernier zones. X axis same as in (A) and (D), Y axis shows parental sequence preservation in Vernier zones. Dashed gray line shows axis between two extremes: Straight CDR graft and Vernier CDR graft. (C) and (F) Overlap of predicted and therapeutic (experimentally validated) humanizing mutations.

To humanize an antibody using Sapiens, we directly leveraged the “recognize and repair” functionality of masked language modeling. First, we fed the Sapiens neural network with a variable region amino acid sequence. Using attention mechanisms, the network recognized non-human residues and output a position-by-residue probability matrix with one row for each position and one column for each of the 20 amino acid types. All possible mutations at all positions were therefore predicted with one pass through the network. We produced the final humanized sequence by taking the most probable predicted residue at each position, except in CDRs, where mutations were ignored and the original parental sequence was preserved. CDR definitions were based on Kabat [37], but IMGT [38], Chothia [39] and North [40] definitions are also supported in BioPhi. Multiple iterations of this process can be performed to humanize the sequence further. After one pass, we refer to a humanized sequence as Sapiens*1, with subsequent passes as Sapiens*2, Sapiens*3, and Sapiens*4.

In the BioPhi web platform, we implemented an integrated pipeline that humanizes sequences using Sapiens (with given number of iterations) or CDR grafting and evaluates their humanness using OASis (Supplementary Figures 7,8). Additional back-mutations or forward mutations to the sequence can then be performed manually by the user through the BioPhi Designer interface, which suggests residues based on Sapiens score, positional residue frequency or nearest germline sequence (Supplementary Figure 9).

We report the humanization results of the 25 antibody pairs in Figure 5. We used OASis identity curves to report on humanness of the humanized sequences across different prevalence thresholds (Figure 5A). As expected, most human-like sequences on average were produced by Straight CDR grafting. The average OASis curve of Sapiens*3 intersected with that of experimental sequences, indicating comparable humanness, followed by Vernier CDR grafting with marginally lower humanness. Interestingly, based on OASis, the Hu-mAb method achieved distinctively lower humanness compared to other methods. Since Hu-mAb sequences were optimized to achieve the same Hu-mAb humanness scores as the experimental sequences, we further investigated this by comparing Hu-mAb scores to T20 and OASis identity scores. When comparing each pair of humanized sequences (Hu-mAb optimized and experimentally validated), both T20 and OASis identity were lower for the sequence that underwent Hu-mAb score optimization, even though they were designed to achieve the same Hu-mAb score (Supplementary Figure 10). The discrepancy suggested that the Hu-mAb metric was no longer an unbiased estimator when applied to Hu-mAb-optimized sequences.

Next, we evaluated the humanness-preservation tradeoff, visualizing the humanness increase compared to the preservation of parental sequence. The increasing Sapiens iterations produced the expected trend of increased humanness and decreased preservation. Sapiens*3 achieved the same humanness as experimental sequences (+34% absolute increase), while achieving higher preservation of the full variable region (86% and 80% respectively, Figure 5B) and same preservation of Vernier zones (86%, Figure 5C).

To provide a breakdown of these average preservation results, we calculated preservation separately at each Kabat position (Supplementary Figure 11). We identified slight differences in preservation of different Vernier positions, where Sapiens*3 achieved higher preservation notably in H48, H67, H69 and H78, while achieving lower preservation notably in H27, H28, H30, H71 and H73.

Finally, we evaluated the humanizing mutation overlap between the sequences produced by automated humanization methods and those validated experimentally (Figure 5D, Supplementary Figure 12,13). For each humanizing mutation to the parental sequence, we determine whether it was shared (made in predicted as well as experimental sequence) or whether it was only made by either one of the methods. Sapiens*1, *2 and *3 achieved highest fractions of shared mutations (humanizing mutation precision, Table 2). To further validate the probabilistic properties of Sapiens predictions, we evaluated the predicted Sapiens score of humanizing mutations and back-mutations made in the experimental sequences. We observed that most mutations achieved the first or second highest Sapiens score (Supplementary Figure 14).

While we report average results, deviation across the 25 antibodies was substantial. To facilitate readers’ closer inspection, we provide the 25 predicted humanized sequences in the Supplemental Information, along with the individual antibody results of the humanness-preservation tradeoff (Supplementary Figure 15) and T20 humanness (Supplementary Figure 16).

Taken together, based on the *in silico* evaluation, BioPhi provides a toolbox of automated humanization methods that are competitive with manual humanization by expert. Moreover, guided by the predictive tools, users can perform further adjustments to the final sequence and produce multiple sequence variants.

### Sapiens rediscovers humanizing mutations of 152 therapeutic antibodies

To evaluate Sapiens and other humanization methods on a larger scale, we reconstructed putative parental sequences of 152 humanized antibody therapeutics. Humanized antibodies generally contain framework regions of human origin and CDRs of parental origin (usually mouse). Therefore, we developed a parental sequence reconstruction strategy based on sequence similarity of CDR regions against all 169,870,516 non-human sequences in OAS, and applied it to 152 humanized sequences from TheraSAbDab [41] (see Methods).

To evaluate the fidelity of the reconstructed parental sequences, we compared these with known parental sequences of 22/152 antibodies which were present in the Hu-mAb 25 pairs dataset. On average, the reconstructed sequences achieved 92% average heavy chain sequence identity and 93% average light chain sequence identity with the known parental sequences. Since each parental sequence was already very similar to the humanized sequence that was used to perform the reconstruction (80% heavy chain identity and 82% light chain identity on average in the 22 pairs), total sequence identity could overestimate the recovery performance. Therefore, we also measured sequence identity based only on mutated positions – positions that were different between the humanized sequence and the parental sequence. In that regard, our strategy correctly recovered 65% mutations in the heavy chain and 72% in the light chain (where a random baseline would recover 5% since there are 20 residues to choose from). Finally, we compared the CDR-based recovery to a strategy based on full sequence homology. Parental sequences recovered that way achieved only 78% and 82% sequence identity with the known heavy chain and light chain sequences respectively, correctly recovering 37% and 66% framework mutations respectively. Therefore, we conducted the following analysis on the CDR-homology-based results.

Similar to the 25 pairs dataset, we used the 152 reconstructed parental sequences as input to each humanization method and evaluated their ability to rediscover the humanized therapeutic sequences. We evaluated each method based on two scenarios. In the first scenario, the humanization method was not instructed with any specific germline and needed to choose it automatically. In the second scenario, the humanization method was instructed with a specific germline gene corresponding to the known humanized therapeutic sequence, simulating a use-case when the germline is requested manually by the user.

With automatic germline selection, each humanization method selected the target germline in a different fashion. In CDR grafting, we selected the germline V and J genes with highest sequence identity to the input sequence. In Hu-mAb, the germline family was selected based on which of the Hu-mAb models achieves highest score on the input sequence. In Sapiens, germline selection was implicit – the neural network was not provided with any germline annotations during training, any germline knowledge was trained directly from the repertoire sequence corpus. Consequently, Sapiens predicted humanizing mutations that maximized the likelihood of seeing the sequence in the training corpus, conditioned on the particular input sequence. Sapiens*1 achieved the same humanness as experimental sequences (+30% absolute increase), while achieving higher preservation of the full variable region (89% and 84% respectively, Figure 6A) as well as in Vernier zones (89% and 87% respectively, Figure 6B). Sapiens*1 also achieved the highest fraction of mutations shared with the experimental sequence (Figure 6C, Table 2). Compared to other methods, Hu-mAb achieved lower overlap due to frequently choosing different germline genes.

With manual germline selection, the target germline gene corresponded to the germline of the humanized therapeutic sequence. In this scenario, the responsibility of the humanization method was reduced mostly to determining which positions in the sequence should be humanized and which should be back-mutated (remain parental), since to a large extent, the residues themselves were already defined based on the chosen germline. In Sapiens and CDR grafting, we provided the germline genes (e.g. IGHV1-46 and IGHJ4 for heavy chain, IGKV1-39 and IGKJ1 for light chain), and determined the allele based on highest sequence identity of each germline sequence with the humanized therapeutic sequence. In Hu-mAb, we provided only the V germline family (e.g. IGHV1 and IGKV1) since more fine-grained selection was not supported. Since Sapiens was trained on all germlines of a given chain type combined, there was no direct way to choose the target germline for Sapiens humanization. To circumvent this issue, we generated Sapiens predictions by first performing Vernier CDR grafting to the target germline, and then applying Sapiens to humanize the sequence further and resolve potential issues at region boundaries. Sapiens*1 achieved higher humanness than experimental sequences (+34% and +30% absolute increase respectively) at the cost of achieving lower preservation of the full variable region (83% and 84% respectively, Figure 6D) as well as in Vernier zones (83% and 87% respectively, Figure 6E). Interestingly, with manual germline assignment, Vernier CDR grafting achieved comparable humanness to experimental sequences (+30% absolute increase) and superior preservation in full variable region (85% and 84% respectively) and in Vernier zones (100% and 87% respectively). Highest mutation overlap with experimental sequences was achieved by Vernier CDR grafting followed by Sapiens*1 (Figure 6F, Table 2). In eight cases, Sapiens*1 predicted a sequence that differed in only one mutation from the experimental sequence, and in one case the sequences were identical (Supplementary Figure 18). This was more than Vernier CDR grafting which produced one case with one mutation difference and one case where the sequence was identical (Supplementary Figure 19).

Using a personal computer with 8 cores, we were able to humanize the 152 antibodies using the BioPhi web interface in 5.2 minutes. We thus concluded that BioPhi was able to humanize sequences at scale while recovering high overlap with therapeutic sequences, with target germlines automatically assigned or manually selected by the user.

## Discussion

We developed BioPhi, an open platform for protein engineering that integrates novel humanization and humanness evaluation methods. The BioPhi automated humanization workflow enables antibody humanization in bulk using a novel humanization method based on deep learning on largescale natural antibody repertoires (Sapiens) or canonical humanization methods based on CDR grafting. *In silico* evaluation demonstrated that the sequences produced by our humanization methods are competitive with those validated experimentally and produced by expert methods. Humanized sequences can further be adjusted manually using the BioPhi Designer functionality. The BioPhi humanness report enables humanness evaluation using a novel method based on 9-mer peptide search (OASis) and traditional methods based on nearest germline sequence identity and positional residue frequency. This enables identifying non-human peptides and residues and suggest viable point mutations based on humanness criteria that are interpretable while exposing the vast sequence diversity of natural antibody repertoires. As an extensible platform that integrates data-driven methods, BioPhi is poised to grow as new datasets become available that continue to connect the two parallel lines of research — adaptive immune repertoire sequencing and antibody engineering.

Established humanness evaluation methods can distinguish between human and non-human sequences, but lack granularity or interpretability. Existing methods based on homology [9][10] are interpretable but lack granularity since they only provide a single score for each chain. Moreover, these underperformed in our analysis compared to more recent approaches such as IgReconstruct [17], which could be attributed to the modest size of their reference sequence libraries. Interestingly, Germline content, a baseline method we implemented based only on percent sequence identity to nearest human germline performed comparably to recent methods while providing both an interpretable and granular score. Nevertheless, antibody germlines provide orders of magnitude smaller sequence space than antibody repertoires, making them more restrictive for antibody engineering applications. Although Hu-mAb [18] has outperformed all other methods including ours by a narrow margin on humanness classification (97.7% and 96.6% ROC AUC respectively for Hu-mAb and OASis medium identity) and on immunogenicity prediction (0.34 and 0.28 R^2^ respectively), we identified two drawbacks. Firstly, Hu-mAb produces only a single score per chain. This could be addressed by providing the user with the change of predicted score upon mutation, as implemented in the Hu-mAb humanization protocol. Secondly, the Hu-mAb score is not interpretable. Although random forest models are robust estimators thanks to their randomized ensemble architecture, this makes their individual predictions difficult to interpret. In contrast, OASis provides a high-accuracy humanness score that is both granular and interpretable, guided by the principles of foreign protein recognition via the processing, display and recognition of peptides.

To be able to compare humanization methods, in this study we mostly reported average performance results across multiple sequences. However, we acknowledge that the deviation in performance across sequences is substantial. Different methods were more successful in different cases, further encouraging the assembly of a diverse arsenal of humanization methods. We imagine this will be enabled by our open-source BioPhi platform.

When evaluating the performance of an automated humanization method, it is crucial to compare it to simple but realistic baselines. Such comparison has not been performed in previous studies [13][18]. In this study, we implemented two baseline methods based on CDR grafting – the Straight CDR graft, which achieved high humanness while preserving parental sequence in Kabat CDRs, and the Vernier CDR graft, which produced less human sequences in exchange for additionally preserving all parental residues in Vernier zones. Both methods performed remarkably well in terms of the overlap with experimentally validated sequences, especially when supplied with the target germline gene.

When humanizing a sequence iteratively by humanness score optimization, the produced sequence should be validated by an independent humanness score, since even small errors in humanness estimation will be amplified during its optimization. We believe this was the cause of inferior OASis and T20 scores for sequences produced by steepest descent optimization in Hu-mAb as compared to results achieved by Sapiens or expert. This further supports our decision to develop separate methods for humanization and humanness evaluation.

Due to its laborious nature, humanization is traditionally performed after the candidate pool has been reduced to a handful of sequences by multiple rounds of binding assays, functional assays and basic biophysical characterization. As these candidate pools are growing with the advent of high-throughput protein production and screening, automated humanization and other antibody engineering methods can help exploring a larger and more diverse sequence space earlier in the process. In the first phase, automated methods can serve as a guide for human-assisted batch humanization and engineering. Ultimately, as their performance improves, they can be combined with other prediction tools to perform holistic *in silico* antibody engineering that will enable humanizing a sequence together with engineering desired properties. In that view, the multiple stages of experimental validation would be used for iterative optimization of the candidate pool on all required properties at once, rather than devising separate stages for affinity optimization, developability optimization, liability mitigation, and de-immunization.

Sapiens is trained with a general goal of recognizing masked or mutated residues and repairing them based on the sequence context. This mechanism could be applied in conjunction with additional optimization criteria to explore vast search spaces of mutations for different antibody engineering tasks. For example, joint optimization of humanness and structural stability prediction was previously used to produce successful humanized candidates [12]. Sapiens could also be used to propose viable point mutations for post-translational modification liability mitigation, both in frameworks and CDRs. More developable molecules could be produced by enriching the Sapiens training set for sequences with properties linked to favorable developability profiles [28] or pairing Sapiens with homology modeling and structure-based developability prediction methods [30].

Since datasets with target measurements are sparse and the input space is enormous, the protein engineering field has started showing interest in unsupervised or self-supervised learning, inspired by the recent progress in natural language processing. By training deep neural networks on large databases of unlabeled sequences, compact numeric representations of proteins can be created, enabling transfer learning on problems with substantially smaller datasets. Notably, this was recently demonstrated in MSA Transformer [25], where residue-residue contact information emerged directly from attention weights after unsupervised learning on multiple sequence alignments. In Sapiens, we have not observed any emergence of such a strong signal, although a pattern of attention from the CDR2 loop to CDR1 and DE loops was clearly present, corresponding to the three-dimensional structure of antibody loops. More work is yet to be done on the representation learning of antibody sequences – curating databases of sequences with favorable properties, optimizing neural network architectures and hyperparameters, and more importantly, inventing large-scale self-supervised tasks that are predictable yet complex enough to force the model to create meaningful inner representations using unlabeled data.

Analogous to natural language processing, antibody humanization and protein engineering methods in general are lacking a single “ground truth”, which makes their *in silico* evaluation and consequently their improvement challenging. Even though in isolation, the humanness-preservation tradeoff achieved by Sapiens is comparable to expert, we understand that further experimental validation is necessary. However, as artificial-intelligence-driven tools such as natural language translation have demonstrated, even before an automated approach achieves human-level performance, it can provide value to the community and create novel opportunities for a new generation of advanced tools and approaches.

## Methods

### OASis peptide database

Unaligned amino acid sequences were obtained in JSON format from unpaired OAS database (accessed Nov 2019). Next, studies with human subject information were selected. Only subjects containing at least 10,000 redundant complete sequences for given chain type were selected, which yielded 118,713,869 sequences from 231 subjects (225 with available heavy chains, 154 with light chains, 148 with both) from 26 studies. For each OAS subject, all overlapping 9-mer peptides were extracted from the amino acid sequences. Heavy chain peptides that appeared only in one subject were removed (corresponding to minimum prevalence of 1% of subjects). An inverted index data structure was created where each distinct peptide points to a list of subjects in which it appears together with the number of occurrences. This was stored along with a subject metadata table in an SQLite database (22GB uncompressed) with an index on the peptide field to speed up querying (less than 1ms per peptide single-threaded).

### Diversity evaluation

Germline sequences were downloaded from IMGT Gene-DB “F+ORF+in-frame P amino acid sequence” for homo sapiens (containing 60 IGHD, 13 IGHJ, 406 IGHV, 9 IGKJ, 108 IGKV, 12 IGLJ and 98 IGLV genes). Overlapping 9-mer peptides were extracted separately from each gene.

A representative sample of OAS sequences for the UMAP visualization was generated by randomly sampling each OAS subject for 25 aligned sequences from each heavy V gene family and 15 aligned sequences from each light V gene family, only complete sequences were considered. This yielded 39,965 variable heavy and 41,845 variable light chain sequences. Germline V gene sequences with IMGT gaps were collected from IMGT Gene-DB. UMAP embedding [33] was generated by precomputing an all-by-all pairwise sequence identity matrix, where only positions shared by both sequences were considered (to handle missing J region in the germline sequences).

### Evaluation of humanness metrics

Therapeutic sequences with species information were downloaded from IMGT mAb DB [35] by querying for all records having an INN request number and IG Receptor Type. OASis identity curves across all prevalence thresholds were calculated for 198 human, 229 humanized, 63 chimeric and 13 murine therapeutics. Statistical significance of differences in OASis medium identity between each group was calculated using two-sided Mann-Whitney U test.

Ability to separate human and non-human sequences was evaluated using a ROC curve. The expected output was 0.0 for negative class (229 humanized, 41 humanized/chimeric, 63 chimeric, 13 mouse, 6 caninized and 3 felinized therapeutics) and 1.0 for positive class (198 human therapeutics), the predicted output was directly the humanness score. Humanness scores of each therapeutic were calculated as averages of the scores of their chains.

Web services were used for Z-score^1^, T20^2^, HumAb ^3^ and IgReconstruct ^4^ (January 2021). AbLSTM was executed only on heavy chain sequences using the pretrained heavy chain models from the code repository^5^. Germline content was calculated by aligning the sequence using IMGT numbering in ANARCI [42] and computing the percent sequence identity with a concatenation of the nearest human V and J gene from IMGT Gene-DB [32]. MG scores were obtained by correspondence with the authors. No implementation of HSC [11] was publicly available, therefore it was not included in the evaluation. Humanness scores and sequences used for evaluation are provided in Supplementary Table 1.

Correlation with clinical immunogenicity was evaluated on a dataset curated in Hu-mAb study [18], sequences were obtained from IMGT mAb DB [35]. Sequence for catumaxomab was not available, therefore only 217/218 therapeutics were included. Explained variance (R^2^) was calculated after transforming each score to the output range using a simple linear regressor. MHC II binding was predicted using netMHCIIpan 3.1 [43]. A peptide was considered binding if it was predicted below 10 percentile in any of DRB1*0101, 0301, 0401, 0701, 0801, 1101, 1301, 1501 (same alleles as in [28]). Reported immunogenicity along with humanness scores and sequences used for evaluation are provided in Supplementary Table 2.

### OASis humanness metric

First, all overlapping 9-mer peptides were extracted from the input antibody and queried against the OASis database using exact match. Next, the human prevalence of each peptide was calculated as number of subjects containing the given peptide (at least once) divided by the total number of subjects for the given chain type. Finally, a single OASis identity score for the input sequence was calculated as the fraction of peptides with prevalence over a user-specified threshold. OASis percentile score was calculated as the percentile of the OASis identity among the 544 therapeutic antibodies collected from IMGT mAb DB. Using a simple benchmark on a personal computer with 8 cores, BioPhi command-line interface was able to evaluate OASis humanness of 1,000 antibodies in 14 minutes.

### Sapiens training corpus

Unaligned variable region amino acid sequences were downloaded from OAS database (accessed Nov 2019). A heavy chain training set was extracted by sampling 20 million unaligned redundant amino acid sequences from all 38 human heavy chain OAS studies from 2011-2017. The training sequences originated from 24% unsorted, 10% IGHA, 1% IGHD, 1% IGHE, 35% IGHG and 30% IGHM isotypes. A validation set was extracted by sampling 20 million sequences from all 5 human heavy chain studies from 2018. The validation sequences originated from 33% unsorted, 16% IGHA, 1% IGHD, 1% IGHE, 20% IGHG and 28% IGHM isotypes. A light chain training set was extracted by taking all 19,054,615 sequences from all 14 human light chain OAS studies from 2011-2017. A validation set was extracted by taking all 33,133,386 sequences from both 2 human light chain OAS studies from 2018. Studies from 2019 were left out to enable future comparison with new methods on an independent test set.

### Sapiens architecture and training procedure

Sapiens was implemented and trained using fairseq [44] and its RoBERTa module [45]. The Transformer encoder contained 4 layers, 8 attention heads, embedding dimensionality of 128, feed forward network embedding dimensionality of 256. Other parameters of the network were based on RoBERTa defaults. In total, the network contained 568,857 tunable weights. Training procedure was based on “masked_lm” training task with 15% masking probability. Label-smoothed cross-entropy with epsilon of 0.1 was used to avoid penalizing the model for making incorrect yet plausible predictions, reflecting the inherent unpredictability of the sequence. A 10% dropout and variable rates of weight decay were used to avoid overfitting. Separate models were trained for the heavy chain and the light chain. The heavy chain model was trained for 700 epochs (166 epochs with learning rate of 1e-4, then further with learning rate of 1e-3) using Adam with default parameters. The light chain model was trained for 300 epochs with learning rate of 1e-04. No hyperparameter tuning was performed. Towards the end of the training procedure, the increase of validation performance started slowing down, but still did not plateau, suggesting that additional training or less conservative regularization techniques could improve performance further.

Although antibody sequences are commonly numbered and aligned for machine learning applications [13][18][26], unaligned sequences were used for three reasons. First, such alignment is only applicable to antibodies and T-cell receptors, so it would render the method inapplicable to other domains in the future. Second, while alignment can help relate conserved positions to each other, it can also conceal motifs found in a particular sequence by fragmenting it with artificial gaps. Thirdly, by using an unaligned sequence, the model was forced to recognize the conserved positions on its own and therefore learn a richer inner representation. Nevertheless, evaluation of alternative training and validation schemes was not performed in this study, so aligned sequence input as well as other subsampling and validation split strategies could also be considered.

### Sapiens attention visualization

Attention weights were collected from 64 heavy chain sequences from IMGT mAb DB that were 120 amino acids in length and were composed of the following AHo positions: 1-7, 9-27, 29-33, 39-61, 65-113, 133-149. Attention weights for each of the 64 sequences were extracted from the Sapiens network, averaged across subjects and attention heads in each layer. An increase in attention to a given region was calculated by comparing mean attention to all positions in the given region compared to mean attention to all positions in other regions. Attention weights for Pembrolizumab heavy chain were visualized in its PDB structure 5B8C using ProVis [46] and nglview [47].

### CDR grafting

IMGT-aligned human germline V and J gene sequences were collected from IMGT Gene-DB. The grafting process consisted of five steps. First, the input sequence was IMGT-aligned using ANARCI. Next, the nearest human V and J gene sequences were selected based on sequence identity (optionally filtered for sequences from specified gene or gene family), and merged into a single sequence with IMGT gaps. Next, the input and germline sequences were renumbered to a user-specified numbering scheme (Kabat by default). In Straight CDR grafting, CDR residues from the input sequence (now based on the renumbered scheme) were inserted at the corresponding positions in the germline sequence. In Vernier CDR grafting, parental Vernier zone residues are grafted along with the CDRs, in other words, these were additional “back-mutations”. Vernier zones were defined based on [8]. Both methods were released in a new open-source package AbNumber^6^.

### Humanization methods evaluation

Validation set of 25 humanized sequences paired with their known parental sequences and Hu-mAb predictions was acquired from the HumAb study [18]. Hu-mAb predictions from the 152 putative parental sequences were generated using the Hu-mAb web server (March 2021). Humanness of predicted sequences was evaluated using OASis and the T20 web server. Humanness change was computed as average absolute difference of OASis medium identity of the humanized sequence and the parental sequence. Parental preservation was calculated as average sequence identity of the parental and humanized sequence under Kabat numbering. To produce Venn diagrams that evaluate humanizing mutation overlap between the experimentally validated sequence and a predicted sequence, all mutations from parent to the experimental or predicted sequence were pooled together and classified into three categories: 1) Shared mutations were made in both the experimental and the humanized sequence (identical residues on the same Kabat position) 2) Experimental only mutations were made only in the experimental sequence 3) Predicted only mutations were made only in the predicted sequence. Finally, humanizing mutation precision was calculated as number of shared mutations divided by total number of predicted mutations. Using a simple benchmark on a personal computer with 8 cores, BioPhi command-line interface was able to humanize 1,000 antibodies in 2.3 minutes (using Sapiens without OASis evaluation).

### Recovering 152 putative parental sequences

Records with “-zumab” suffix were collected from TheraSAbDab, totaling 164 humanized therapeutics. To estimate which positions (apart from CDRs) in the humanized therapeutic sequences came from their parental sequence, framework residues with <1% positional frequency were identified. The frequency was calculated based on a subset of 4 million human OAS sequences created by sampling 10,000 complete sequences from each OAS subject. On average there were 1.3 rare framework residues in the heavy chain and 1.0 in the light chain.

Each humanized sequence was IMGT-aligned using ANARCI and compared against all 169,870,516 non-human IMGT-aligned sequences from OAS (94.3% from mouse, 2.5% from rat, 1.8% from rabbit, 0.8% from rhesus and 0.7% from camel). The putative parental sequence template was selected based on highest sequence identity in CDRs. In case multiple sequences with same CDR identity were found, the one with highest framework identity was selected. To preserve only high-confidence matches, therapeutics with less than 60% CDR identity with the nearest match in heavy or light chain were discarded, yielding 152 final pairs. The final putative parental sequence was assembled by grafting CDRs and the identified non-human residues of the humanized sequence into the parental OAS hit, essentially performing reverse CDR grafting. This produced a fully non-human sequence with CDRs and other non-human residues from the humanized therapeutic and frameworks from the non-human OAS hit. For 22/152 therapeutics, known parental sequences were obtained from [18]. Sequence identity was calculated as percentage of identical residues in the known and recovered parental sequence using Kabat numbering. Recovery mutation accuracy was calculated as number of framework positions that agreed between the two parental sequences while being mutated in the humanized sequence, divided by the total number of mutated positions. Putative parental sequences are provided in Supplementary Table 3.

## Supporting information

Supplementary 25 Antibody Pairs

## Availability

BioPhi web application is available at https://biophi.dichlab.org

BioPhi code repository is available at https://github.com/Merck/BioPhi

The code and data supporting this analysis are available at https://github.com/Merck/BioPhi-2021-publication and in the Supplementary Information.

## Acknowledgements

We thank Jens Christensen, Jacob Lustig, Vincent Antonnuci and Carol A. Rohl for supporting this work. We thank Petr Mejzlik and Geoffrey D. Hannigan for reviewing the final draft of the manuscript.

## Author contributions

DP with the supervision of DS and DB conceived and designed the study. DB assembled the team. JM proposed using 9-mer peptides from OAS for humanness evaluation, DP conceived, implemented and evaluated the OASis identity approach. DP conceived and implemented the Sapiens method and its evaluation, DB consulted the possible applications. DP implemented the BioPhi application and the relevant humanization, humanness evaluation and designer modules. AW, VJ, DS and LFD advised throughout the study. DP wrote the manuscript, DB, DS, VJ, LD and AW edited and contributed to the final draft. All authors reviewed and approved the final draft.

## Footnotes

1 http://www.bioinf.org.uk/abs/shab/

2 https://dm.lakepharma.com/bioinformatics

3 http://opig.stats.ox.ac.uk/webapps/humab

4 http://meilerlab.org/index.php/servers/IgReconstruct

5 https://github.com/vkola-lab/peds2019

6 https://github.com/prihoda/AbNumber

