## Supplementary Figures for "BioPhi: A platform for antibody design, humanization and humanness evaluation based on natural antibody repertoires and deep learning"

| | Antibody | | | Heavy chain | | | | Light chain | | | |
|---|---|---|---|---|---|---|---|---|---|---|---|
| Name | OASis Percentile | OASis Identity | Germline Content | OASis Percentile | OASis Identity | Germline Gene | Germline Content | OASis Percentile | OASis Identity | Germline Gene | Germline Content |
| Golimumab | 93% | 96% | 92% | 91% | 93% | IGHV3-30*01 | 89% | 100% | 100% | IGKV3-11*01 | 96% |
| Muromonab-CD3 | 0% | 37% | 66% | 8% | 51% | IGHV1-46*01 | 69% | 0% | 21% | IGKV3-11*01 | 62% |
| Pembrolizumab | 26% | 72% | 79% | 28% | 69% | IGHV1-2*02 | 76% | 32% | 76% | IGKV3-11*01 | 82% |

**Supplementary Figure 1:** BioPhi bulk antibody humanness evaluation report. Humanness is evaluated using OASis percentile, OASis identity and nearest human germline identity.

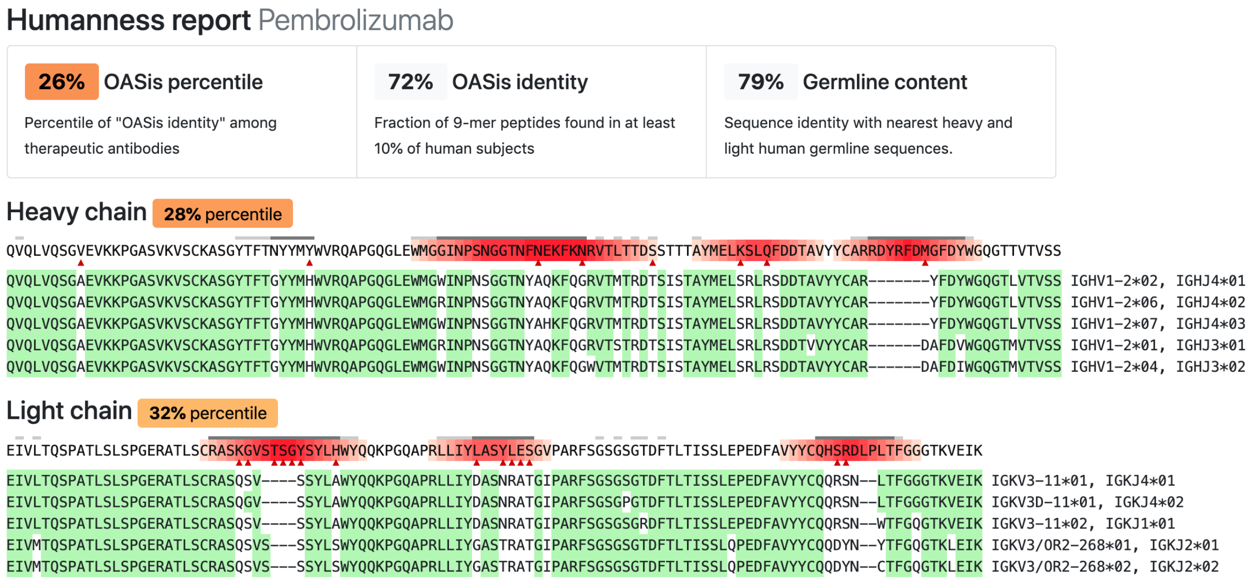


**Supplementary Figure 2:** BioPhi humanness evaluation report of Pembrolizumab. Humanness of each residue is annotated using red gradient based on the number of non-human overlapping 9-mer peptides at given position (peptides that don't satisfy the prevalence threshold). Rare residues at given position are marked with red triangles. Each chain is aligned to five nearest V and J germline gene sequences.

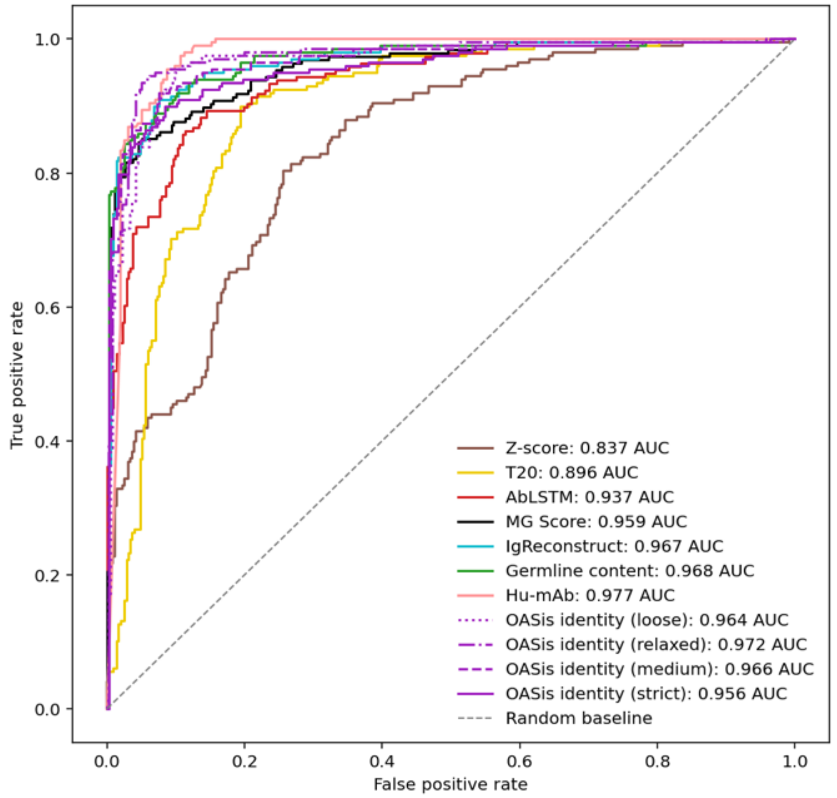


**Supplementary Figure 3:** Classification performance of humanness scores to distinguish between sequences that are human and sequences that are non-human (humanized or other species) visualized using a ROC curve.

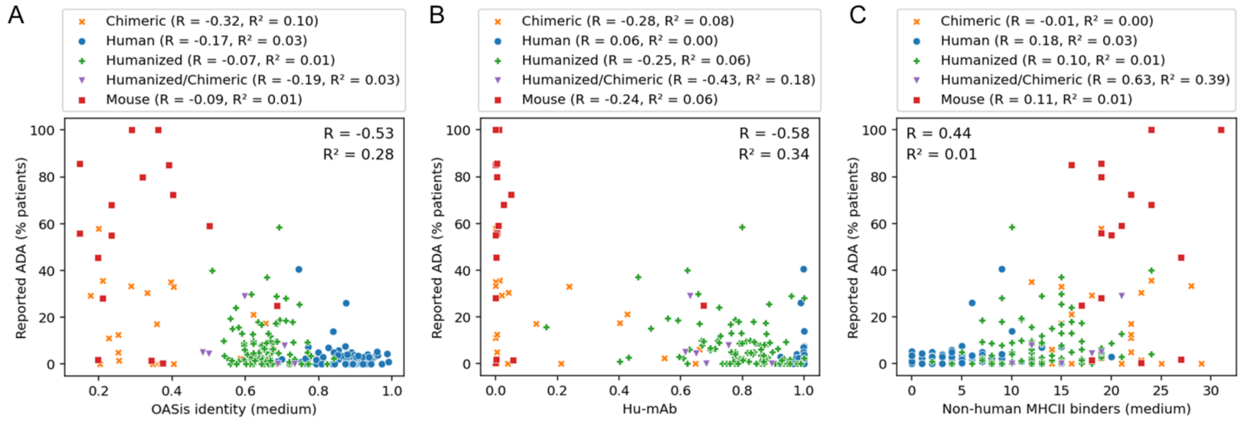


**Supplementary Figure 4:** Relationship of humanness scores and reported immunogenic anti-drug antibody (ADA) response of therapeutic antibodies. X axis shows the humanness score (OASis identity, Hu-mAb and Non-human MHC-II binders in A, B and C respectively), Y axis shows reported percentage of patients with observed anti-drug antibody response. Although the correlation with immunogenicity is visible across all humanness scores, this signal appears to be driven largely by the species of origin, therefore R and $R^2$ results for each individual species are also reported in the figure legends.

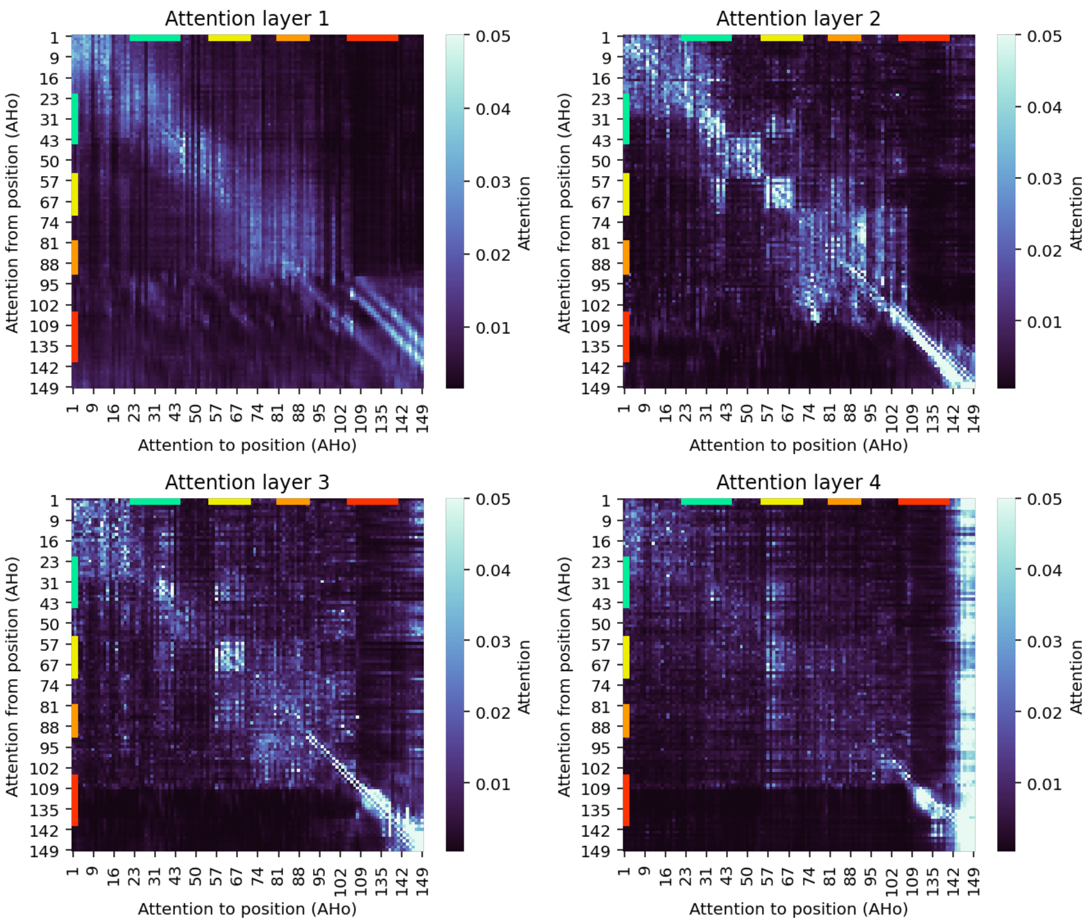


**Supplementary Figure 5:** Average attention matrices in the four layers of the Sapiens neural network. Average attention was calculated on 64 heavy chain sequences from IMGT mAb DB that were composed of the same positions under Aho numbering: 1-7, 9-27, 29-33, 39-61, 65-113, 133-149.

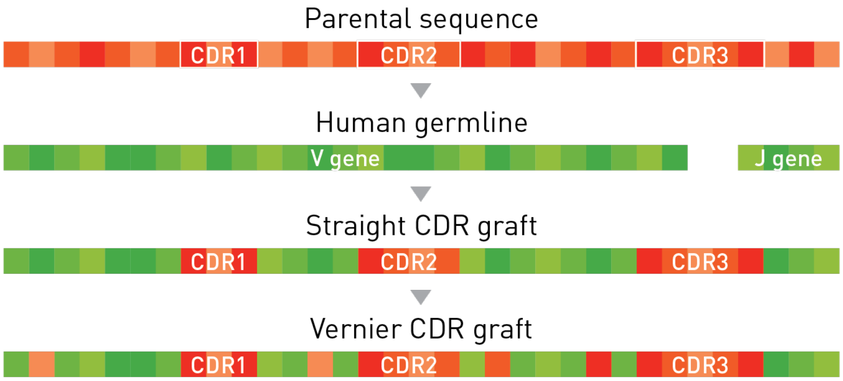


**Supplementary Figure 6:** Simplified illustration of the CDR grafting procedure. The Straight CDR graft was created by inserting Kabat CDR regions into nearest human germline V and J genes. The *Vernier* CDR graft was created from the Straight CDR graft by additionally back-mutating all Vernier zone positions to the parental residues.

| | OASis Percentile | | OASis Identity | | Germline identity | | Germlines | | Humanizing mutations | |
|---|---|---|---|---|---|---|---|---|---|---|
| Name | Before | After | Before | After | Before | After | VH | VL | VH | VL |
| AntiCD28 | 6% | 19% | 52% | 70% | 66% | 75% | IGHV4-4*08 | IGKV4-1*01 | 13 | 17 |
| Bevacizumab | 4% | 50% | 50% | 79% | 71% | 84% | IGHV7-4-1*02 | IGKV1-33*01 | 15 | 15 |
| Campath | 9% | 50% | 62% | 79% | 72% | 84% | IGHV3-71*01 | IGKV1-39*01 | 16 | 10 |
| Herceptin | 3% | 20% | 48% | 70% | 63% | 75% | IGHV1-2*06 | IGKV1-33*01 | 14 | 16 |
| Omalizumab | 1% | 30% | 44% | 73% | 67% | 78% | IGHV4-38-2*01 | IGKV4-1*01 | 16 | 14 |

**Supplementary Figure 7:** BioPhi bulk humanization result using Sapiens on five parental sequences from the 25 pairs dataset.

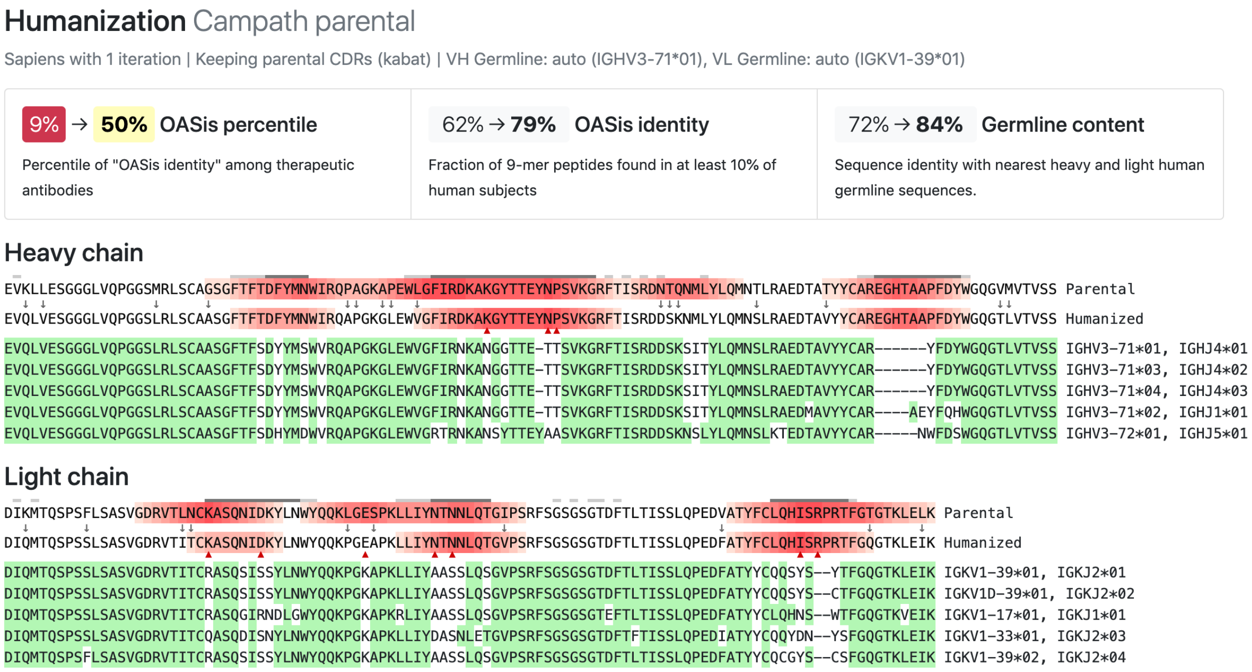


**Supplementary Figure 8** BioPhi Humanization report, humanizing the parental sequence of Campath using Sapiens.

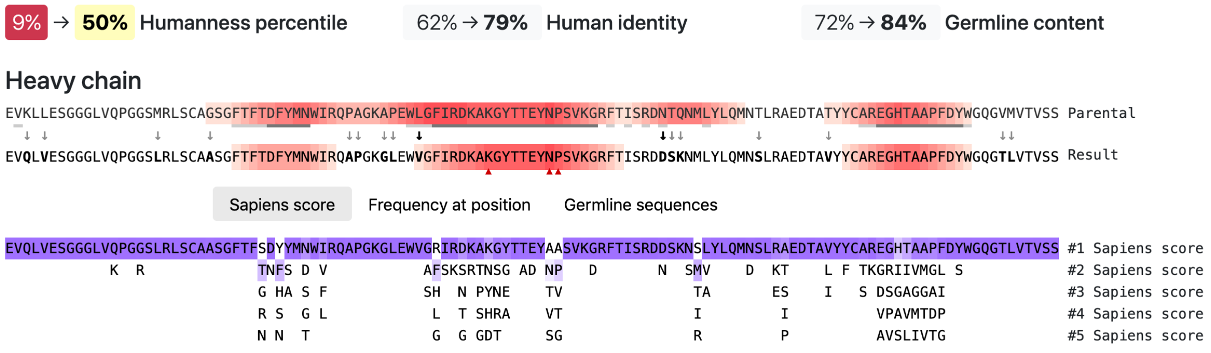


**Supplementary Figure 9:** BioPhi Antibody Designer functionality enabling manual adjustments to a sequence produced by automated humanization (here, 1 Sapiens iteration on Campath parental sequence). Clicking on any residue performs the given mutation to the result sequence. Residues are suggested based on parental sequence, Sapiens score, frequency at position and nearest germline sequences. Any mutation can be performed by clicking at each residue in the result sequence.

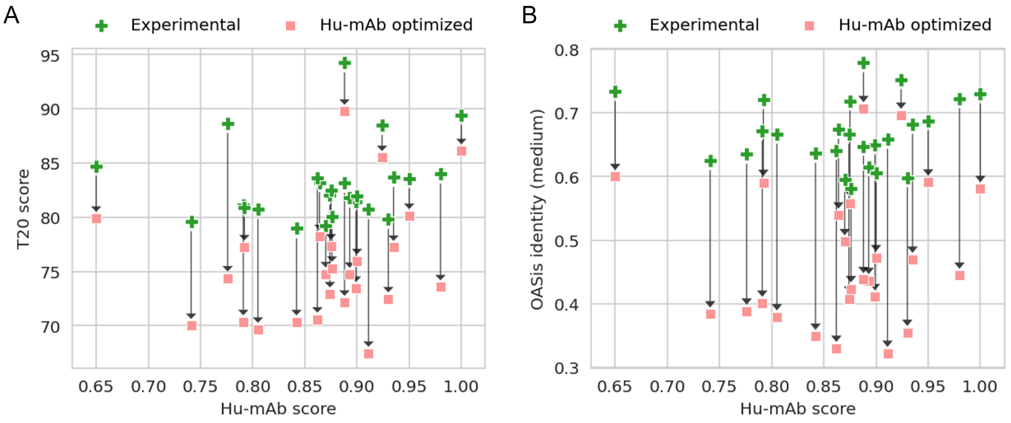


**Supplementary Figure 10:** Comparison of humanness of 25 sequences humanized by Hu-mAb (pink) and humanized experimentally (green) as measured by T20 score (A) and OASis medium identity (B). While each pair of sequences has the same Hu-mAb score, the T20 score and OASis identity scores are consistently lower for the sequences humanized by Hu-mAb (T20=75.6, OASis=47.1 on average) compared to the Experimental sequence (T20=83.1, OASis=66.8 on average).

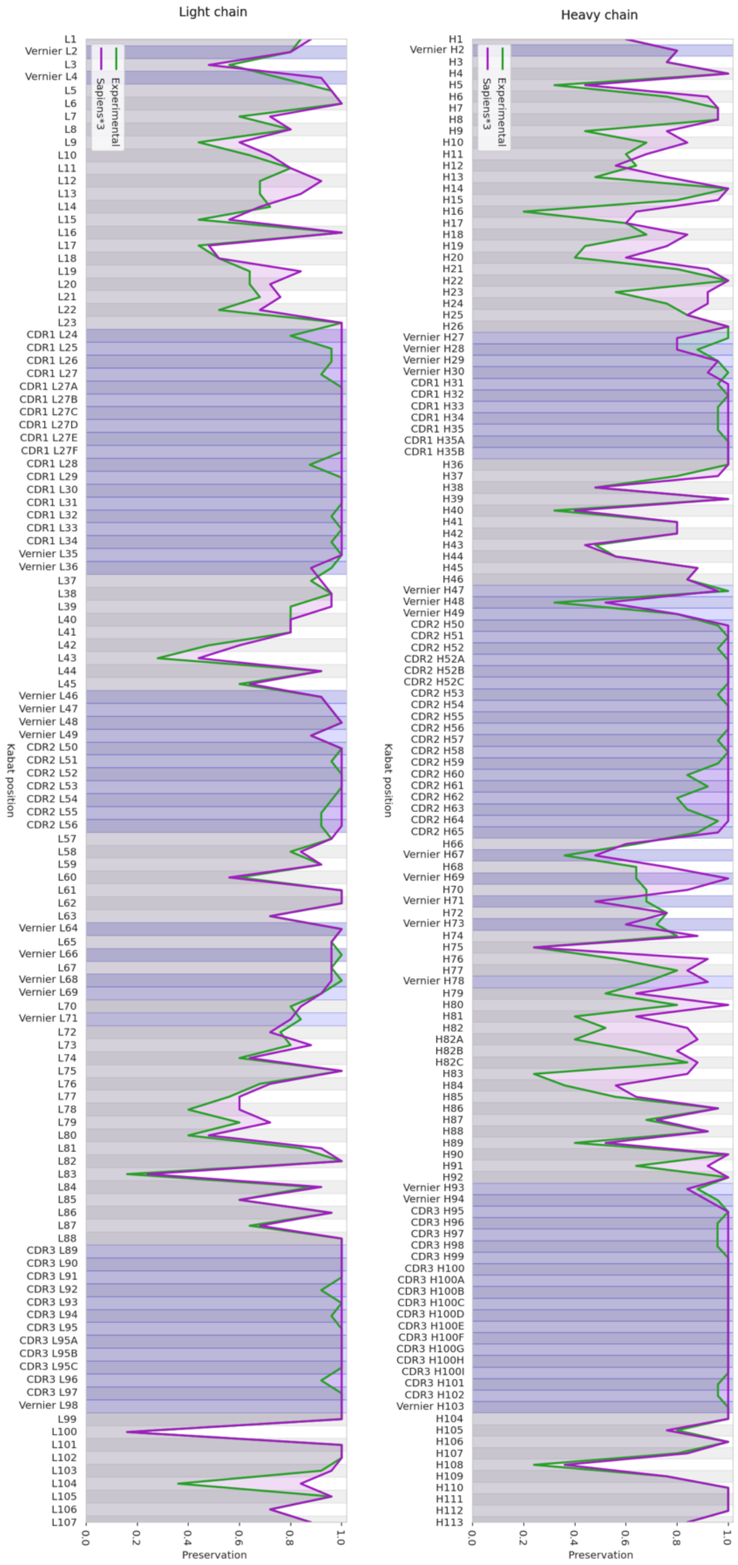


**Supplementary Figure 11:** Fraction of preserved residues between the parental and humanized sequence at each Kabat position. Average results were calculated on the 25 pairs benchmark, sequences humanized experimentally (green) and by Sapiens (purple). Blue regions are highlighting Vernier zones and CDRs.

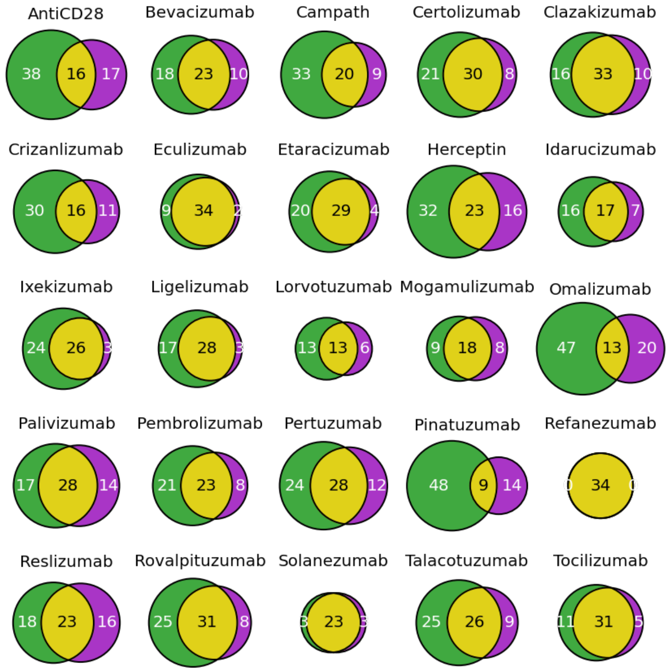


**Supplementary Figure 12:** Overlap of humanizing mutations between Sapiens*3 (purple) and expert (green) for each humanized antibody in the 25 pairs benchmark.

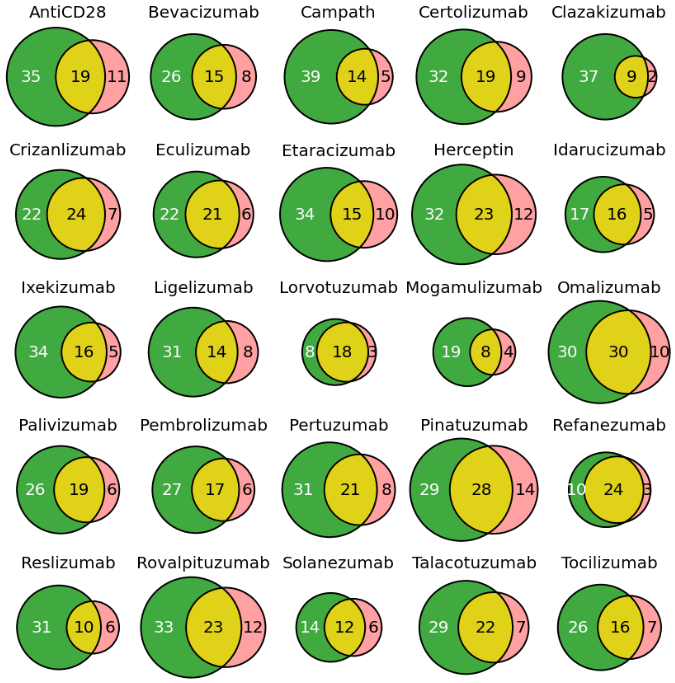


**Supplementary Figure 13:** Overlap of humanizing mutations between Hu-mAb (pink) and expert (green) for each humanized antibody in the 25 pairs benchmark.

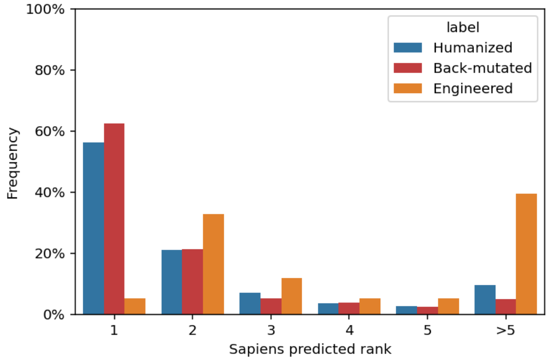


**Supplementary Figure 14:** Predicted Sapiens residue score of humanizing mutations and back-mutations made by 25 experimentally validated humanized sequences. Mutations are further subdivided into three categories: Mutations that match nearest human germline but do not match parent (Humanized), mutations that do not match nearest human germline, but match parent (Back-mutated), mutations that do not match nearest human germline and do not match parent (Engineered). Mutations that match nearest human germline and also match parent (Conserved) are trivial so these were not included in the analysis.

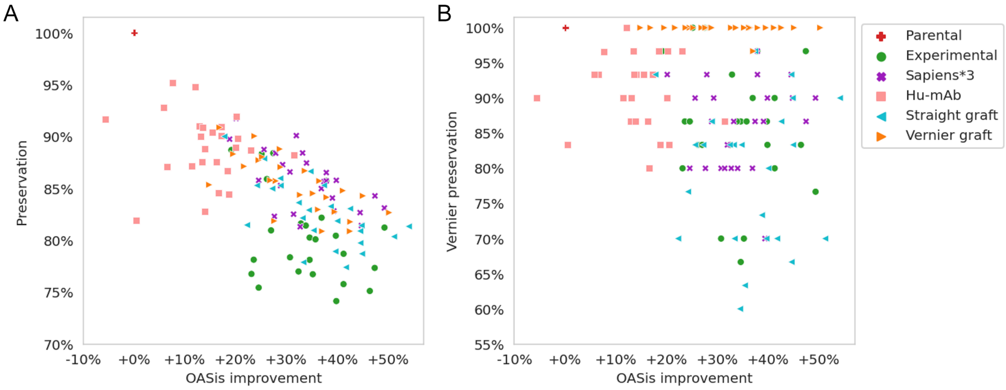


**Supplementary Figure 15:** Humanness-preservation tradeoff for each individual sequence in the 25 pairs benchmark. Preservation shown in full variable region (A) and in vernier zones (B).

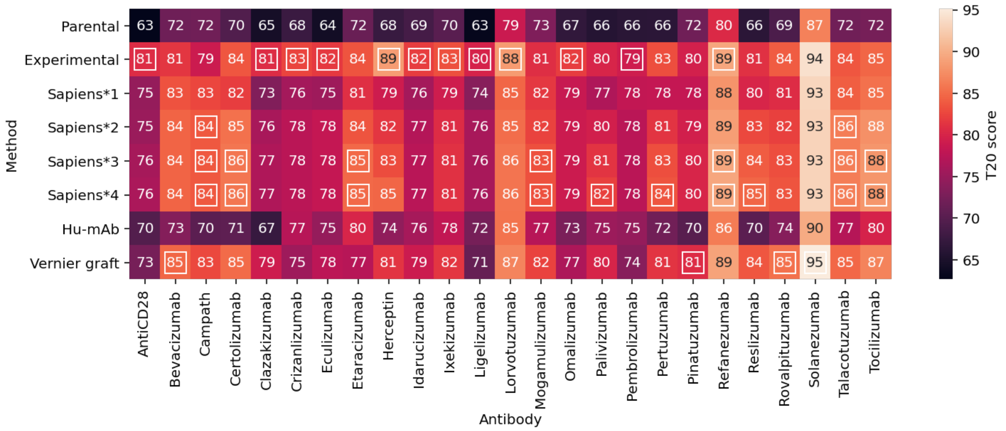


**Supplementary Figure 16:** T20 humanness scores of each individual sequence in the 25 pairs benchmark. Highest value in each column are highlighted with white outline. Results for Straight graft are not included due to their low preservation compared to other methods.

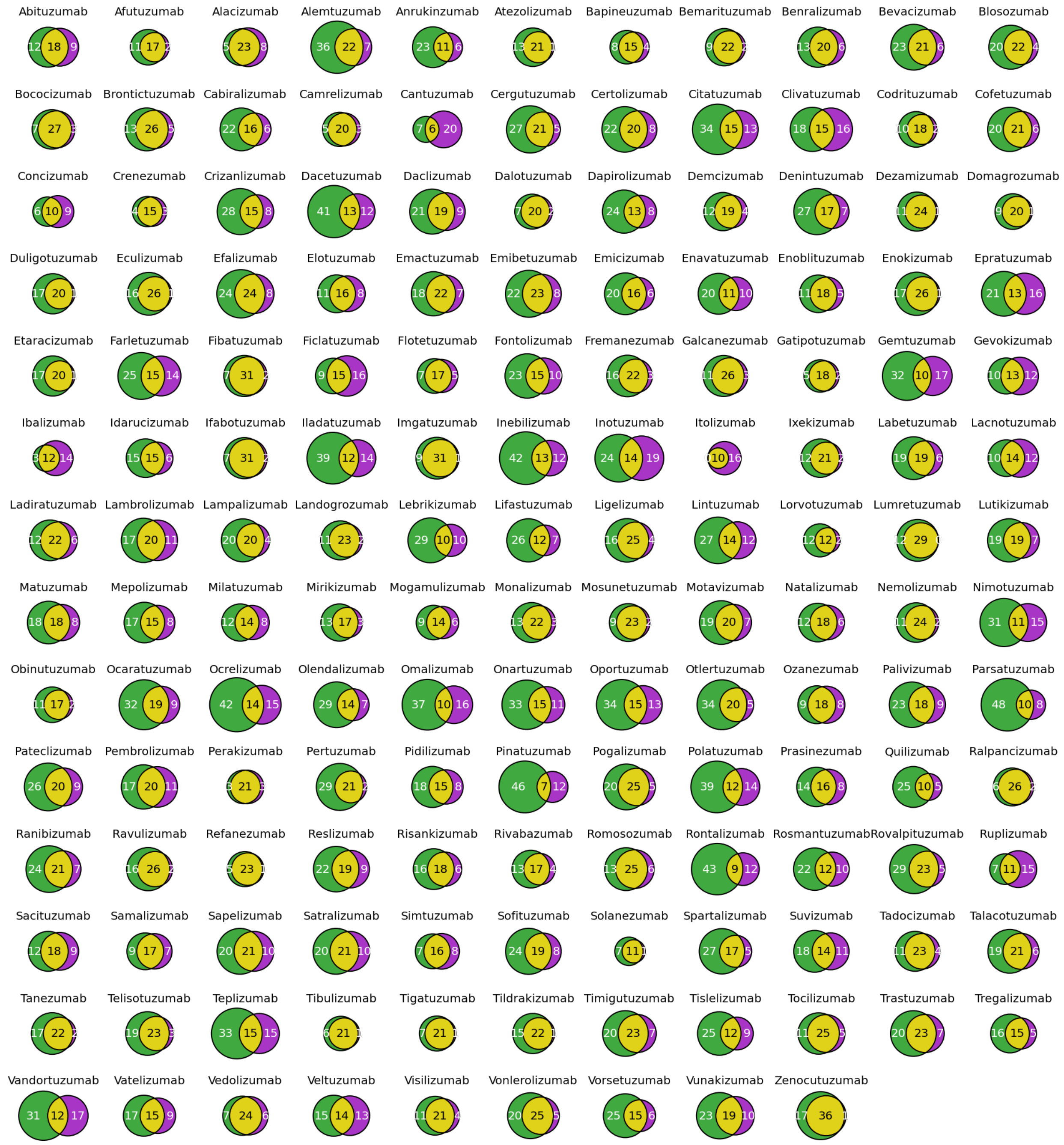


**Supplementary Figure 17:** Overlap of humanizing mutations between Sapiens*1 (purple) and expert (green) for each humanized antibody in the 152 pairs benchmark with automatic germline assignment.

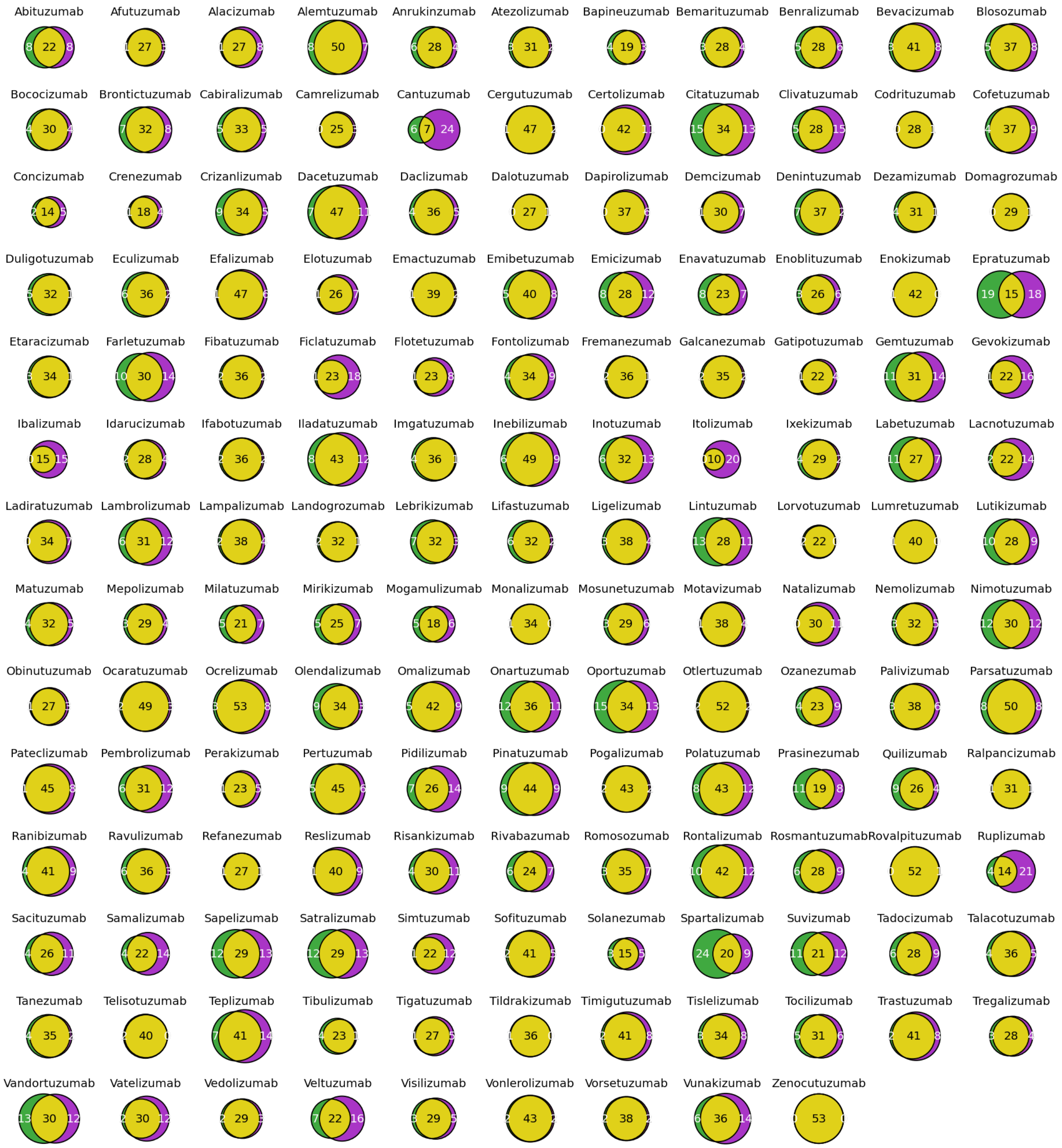

**Supplementary Figure 18:** Overlap of humanizing mutations between Sapiens*1 (purple) and expert (green) for each humanized antibody in the 152 pairs benchmark with manual germline assignment. Eight sequences differ only by one mutation (Codrituzumab, Dalotuzumab, Domagrozumab, Enokizumab, Lumretuzumab, Monalizumab, Rovalpituzumab, Tildrakizumab), one sequence is identical (Zenocutuzumab).

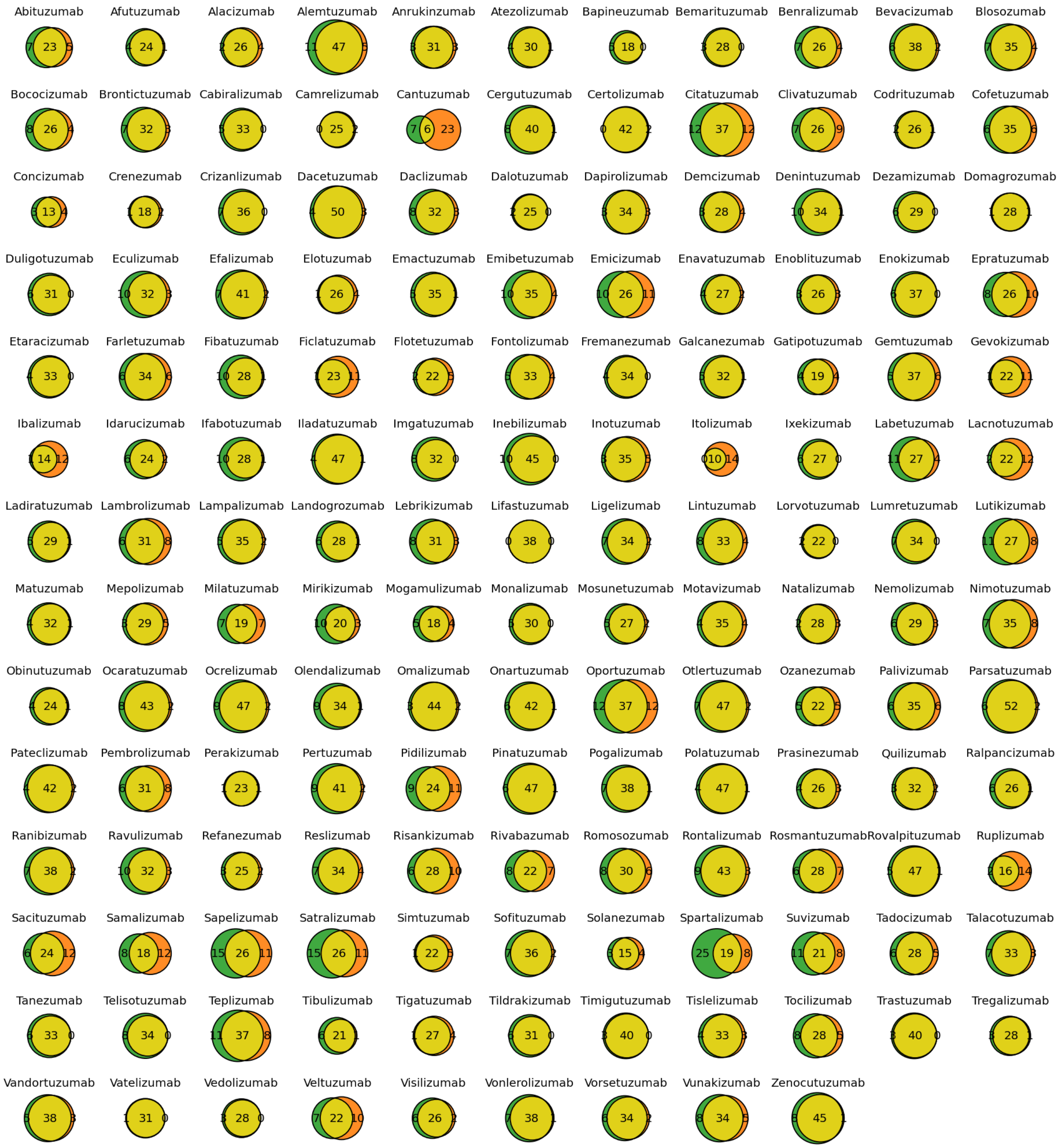

**Supplementary Figure 19**: Overlap of humanizing mutations between Vernier CDR grafting (orange) and expert (green) for each humanized antibody in the 152 pairs benchmark with manual germline assignment. Vatelizumab differs just by one mutation, Lifastuzumab is identical.
