## Supplementary 25 Antibody Pairs for "BioPhi: A platform for antibody design, humanization and humanness evaluation based on natural antibody repertoires and deep learning"

#### Humanization results: 25 sequence pairs

Legend:

- Line 1: Nearest human germline to experimentally humanized sequence
- Line 2: Experimentally humanized sequence
- Line 3: Parental sequence
- Line 4: Sequence humanized by Sapiens (3 iterations)
- Line 5: Nearest human germline to Sapiens humanized sequence

Parental and humanized sequences are colored in red gradient based on number of non-human overlapping 9-mers at given position (based on OASis relaxed identity)

Germline sequences are colored in green when the humanized sequence matches germline

CDR and Vernier regions are shown using dark and light grey border around parental sequence

#### AntiCD28

##### Heavy chain

```
EVQLVQSGGGLVQPGGSLRLSCAGSGFTFSSYAMHWVRQAPGKGLEWVSAIGTGGGTYYADSVKGRFTISRDNAKNSLYLQMNSLRAEDMAVYYCAR--------YFDYWGQGTLVTVSS IGHV3/OR16-10*02, IGHJ4*01
EVQLVQSGGGLVQPGGSLRLSCAGSGFTFSDYGVHWVRQAPGKGLEWVSAIWAGGGTNYASSVMGRFTISRDNAKNSLYLQMNSLRAEDMAVYYCARDKGYSYYYSMDYWGQGTLVTVSS Humanized (Experimental)
EVKLQQSGPGLVTPSQSLSITCTVSGFSLSDYGVHWVRQSPGQGLEWLGVIWAGGGTNYNSALMSRKSISKDNSKSQVFLKMNSLQADDTAVYYCARDKGYSYYYSMDYWGQGTSVTVSS Parental
QVQLQESGPGLVKPSQTLSLTCTVSGGSLSDYGVHWVRQSPGKGLEWLGVIWAGGGTNYNSALMSRVSISLDTSKSQVSLKLNSLTAADTAVYYCARDKGYSYYYSMDYWGQGTTVTVSS Humanized (Sapiens^3)
QVQLQESGPGLVKPSETLSLTCTVSGGSISSYYWSWIRQPPGKGLEWIGYIYTSGSTNYNPSLKSRVTISVDTSKNQFSLKLSSVTAADTAVYYCAR-----YYYGMDVWGQGTTVTVSS IGHV4-4*08, IGHJ6*01
```

##### Light chain

```
DIVMTQSPDSLAVSLGERATINCKSSQSVLYNNKNYLAWYQQKPGQPPKLLIYWASTRESGVPDRFSGSGSGTDFTLTISSLQAEDVAVYYCQQYYS---YTFGQGTKLEIK IGKV4-1*01, IGKJ2*01
DIVMTQSPDSLAVSLGERATINCRASESVEYYVTSLMAWYQQKPGQPPKLLIYAASNVESGVPDRFSGSGSGTNFSLTISSLQAEDVAVYYCQQSRKYVPYTFGQGTKLEIK Humanized (Experimental)
DIETLQSPASLAVSLGQRATISCRASESVEYYVTSLMQWYQQKPGQPPKLLIFAASNVESGVPARFSGSGSGTNFSLNIHPVDEDDVAMYFCQQSRKYVPYTFGGGTKLEIK Parental
DIVMTQSPASLAVSPGERATISCRASESVEYYVTSLMQWYQQKPGQPPKLLIYAASNVESGVPDRFSGSGSGTDFTLTISRVQADDVAVYYCQQSRKYVPYTFGQGTKLEIK Humanized (Sapiens^3)
DIVMTQSPDSLAVSLGERATINCKSSQSVLYNNKNYLAWYQQKPGQPPKLLIYWASTRESGVPDRFSGSGSGTDFTLTISSLQAEDVAVYYCQQYYS---YTFGQGTKLEIK IGKV4-1*01, IGKJ2*01
```

#### Bevacizumab

##### Heavy chain

```
EVQLVESGGGLVQPGGSLRLSCAASGFTFSSYAMSWVRQAPGKGLEWVSAISGSGGSTYYADSVKGRFTISRDNSKNTLYLQMNSLRAEDTAVYYCAK--------YWYFDLWGRGTLVTVSS IGHV3-23*04, IGHJ2*01
EVQLVESGGGLVQPGGSLRLSCAASGYTFTNYGMNWVRQAPGKGLEWVGWINTYTGEPTYAADFKRRFTFSLDTSKSTAYLQMNSLRAEDTAVYYCAKYPHYYGSSHWYFDVWGQGTLVTVSS Humanized (Experimental)
EIQLVQSGPELKQPGETVRISCKASGYTFTNYGMNWVKQAPGKGLKWMGWINTYTGEPTYAADFKRRFTFSLETSASTAYLQISNLKNDDTATYFCAKYPHYYGSSHWYFDVWGAGTTVTVSS Parental
QVQLVQSGSELKKPGASVRISCKASGYTFTNYGMNWVRQAPGQGLEWMGWINTYTGEPTYAADFKRRFVFSLDTSVSTAYLQISNLKADDTAVYFCARYPHYYGSSHWYFDVWGRGTLVTVSS Humanized (Sapiens^3)
QVQLVQSGSELKKPGASVKVSCKASGYTFTSYAMNWVRQAPGQGLEWMGWINTNTGNPTYAQGFTGRFVFSLDTSVSTAYLQISSLKAEDTAVYYCAR--------YWYFDLWGRGTLVTVSS IGHV7-4-1*02, IGHJ2*01
```

##### Light chain

```
DIQMTQSPSSLSASVGDRVTITCRASQGISNYLAWFQQKPGKAPKSLIYAASSLQSGVPSRFSGSGSGTDFTLTISSLQPEDFATYYCQQYNS--WTFGQGTKVEIK IGKV1-16*01, IGKJ1*01
DIQMTQSPSSLSASVGDRVTITCSASQDISNYLNWYQQKPGKAPKVLIYFTSSLHSGVPSRFSGSGSGTDFTLTISSLQPEDFATYYCQQYSTVPWTFGQGTKVEIK Humanized (Experimental)
DIQMTQTTSSLSASLGDRVIISCSASQDISNYLNWYQQKPDGTVKVLIYFTSSLHSGVPSRFSGSGSGTDYSLTISNLEPEDIATYYCQQYSTVPWTFGGGTKLEIK Parental
DIQMTQSPSSLSASVGDRVTITCSASQDISNYLNWYQQKPGKAPKLLIYFTSSLHSGVPSRFSGSGSGTDFTLTISSLQPEDIATYYCQQYSTVPWTFGQGTKVEIK Humanized (Sapiens^3)
DIQMTQSPSSLSASVGDRVTITCQASQDISNYLNWYQQKPGKAPKLLIYDASNLETGVPSRFSGSGSGTDFTFTISSLQPEDIATYYCQQYDN--WTFGQGTKVEIK IGKV1-33*01, IGKJ1*01
```

### Campath

#### Heavy chain

```
QVQLQESGPGLVKPSQTLSLTCTVSGGSIGDYYWSWIRQPPGKGLEWIGYIYYS---GSTYYNPSLKSRVTISVDTSKNQFSLKLSSVTAADTAVYYCAR------YFDYWGQGTLVTVSS IGHV4-30-4*01, IGHJ4*01
QVQLQESGPGLVRPSQTLSLTCTVSGFTFTDFYMNWVRQPPGRGLEWIGFIRDKAKGYTTEYNPSVKGRVTMLVDTSKNQFSLRLSSVTAADTAVYYCAREGHTAAPFDYWGQGSLVTVSS Humanized (Experimental)
EVKLLESGGGLVQPGGSMRLSCAGSGFTFTDFYMNWIRQPAGKAPEWLGFIRDKAKGYTTEYNPSVKGRFTISRDNTQNMLYLQMNTLRAEDTATYYCAREGHTAAPFDYWGQGVMVTVSS Parental
EVQLVESGGGLVQPGGSLRLSCAASGFTFSDFYMNWIRQAPGKGLEWVGFIRDKAKGYTTEYNPSVKGRFTISRDDSKNSLYLQMNSLRAEDTAVYYCAREGHTAAPFDYWGQGTLVTVSS Humanized (Sapiens^3)
EVQLVESGGGLVQPGGSLRLSCAASGFTFSDYYMSWVRQAPGKGLEWVGFIRNKANGGTTE-TTSVKGRFTISRDDSKSITYLQMNSLRAEDTAVYYCAR------YFDYWGQGTLVTVSS IGHV3-71*01, IGHJ4*01
```

#### Light chain

```
DIQMTQSPSSLSASVGDRVTITCQASQDISNYLNWYQQKPGKAPKLLIYDASNLETGVPSRFSGSGSGTDFTFTISSLQPEDIATYYCQQYDN--WTFGQGTKVEIK IGKV1-33*01, IGKJ1*01
DIQMTQSPSSLSASVGDRVTITCKASQNIDKYLNWYQQKPGKAPKLLIYNTNNLQTGVPSRFSGSGSGTDFTFTISSLQPEDIATYYCLQHISRPRTFGQGTKVEIK Humanized (Experimental)
DIKMTQSPSFLSASVGDRVTLNCKASQNIDKYLNWYQQKLGESPKLLIYNTNNLQTGIPSRFSGSGSGTDFTLTISSLQPEDVATYFCLQHISRPRTFGTGTKLELK Parental
DIQMTQSPSSLSASVGDRVTITCKASQNIDKYLNWYQQKPGKAPKLLIYNTNNLQTGVPSRFSGSGSGTDFTLTISSLQPEDFATYFCLQHISRPRTFGQGTKLEIK Humanized (Sapiens^3)
DIQMTQSPSSLSASVGDRVTITCRASQSISSYLNWYQQKPGKAPKLLIYAASSLQSGVPSRFSGSGSGTDFTLTISSLQPEDFATYYCQQSYS--YTFGQGTKLEIK IGKV1-39*01, IGKJ2*01
```

### Certolizumab

#### Heavy chain

```
EVQLVESGGGLVQPGGSLRLSCAASGFTFSSYAMSWVRQAPGKGLEWVSAISGSGGSTYYADSVKGRFTISRDNSKNTLYLQMNSLRAEDTAVYYCAK-----YFDYWGQGTLVTVSS IGHV3-23*04, IGHJ4*01
EVQLVESGGGLVQPGGSLRLSCAASGYVFTDYGMNWVRQAPGKGLEWMGWINTYIGEPIYADSVKGRFTFSLDTSKSTAYLQMNSLRAEDTAVYYCARGYRSYAMDYWGQGTLVTVSS Humanized (Experimental)
QIQLVQSGPELKKPGETVKISCKASGYVFTDYGMNWVKQAPGKAFKWMGWINTYIGEPIYVDDFKGRFAFSLETSASTAFLQINNLKNEDTATYFCARGYRSYAMDYWGQGTSVTVSS Parental
QVQLVQSGSELKKPGASVKISCKASGYTFTDYGMNWVRQAPGQGLEWMGWINTYIGEPIYVDDFKGRFVFSLDTSVSTAYLQINNLKAEDTAVYFCARGYRSYAMDYWGQGTLVTVSS Humanized (Sapiens^3)
QVQLVQSGSELKKPGASVKVSCKASGYTFTSYAMNWVRQAPGQGLEWMGWINTNTGNPTYAQGFTGRFVFSLDTSVSTAYLQICSLKAEDTAVYYCAR-----YFDYWGQGTLVTVSS IGHV7-4-1*01, IGHJ4*01
```

#### Light chain

```
DIQMTQSPSSLSASVGDRVTITCRASQGISNYLAWFQQKPGKAPKSLIYAASSLQSGVPSRFSGSGSGTDFTLTISSLQPEDFATYYCQQYNS--WTFGQGTKVEIK IGKV1-16*01, IGKJ1*01
DIQMTQSPSSLSASVGDRVTITCKASQNVGTNVAWYQQKPGKAPKALIYSASFLYSGVPYRFSGSGSGTDFTLTISSLQPEDFATYYCQQYNIYPLTFGQGTKVEIK Humanized (Experimental)
DIVMTQSQKFMSTSVGDRVSVTCKASQNVGTNVAWYQQKPGQSPKALIYSASFLYSGVPYRFTGSGSGTDFTLTISTVQSEDLAEYFCQQYNIYPLTFGAGTKLELK Parental
DIQMTQSPSSLSASVGDRVTITCKASQNVGTNVAWYQQKPGKAPKLLIYSASFLYSGVPSRFSGSGSGTDFTLTISSLQPEDFATYYCQQYNIYPLTFGQGTKLEIK Humanized (Sapiens^3)
DIQMTQSPSSVSASVGDRVTITCRASQGISSWLAWYQQKPGKAPKLLIYAASSLQSGVPSRFSGSGSGTDFTLTISSLQPEDFATYYCQQANS--YTFGQGTKLEIK IGKV1-12*01, IGKJ2*01
```

### Clazakizumab

#### Heavy chain

```
EVQLVESGGGLVQPGGSLRLSCAASGFTVSSNYMSWVRQAPGKGLEWVSVIYSGGSTYYADSVKGRFTISRDNSKNTLYLQMNSLRAEDTAVYYCAR------AEYFQHWGQGTLVTVSS IGHV3-66*01, IGHJ1*01
EVQLVESGGGLVQPGGSLRLSCAASGFSLSNYYVTWVRQAPGKGLEWVGIIYGSDETAYATSAIGRFTISRDNSKNTLYLQMNSLRAEDTAVYYCARDDSSDWDAKFNLWGQGTLVTVSS Humanized (Experimental)
QSLEESGGRLVTPGTPLTLTCTASGFSLSNYYVTWVRQAPGKGLEWIGIIYGSDETAYATWAIGRFTISKTSTTVDLKMTSLTAADTATYFCARDDSSDWDAKFNLWGQGTLVTVSS Parental
VQLEESGGGLVQPGESLRLSCTASGFSLSNYYVTWVRQAPGKGLEWISIIYGSDETAYATWAIGTISIDKSKNTVYLKMTSLTAADTAVYFCARDDSSDWDAKFNLWGQGTLVTVSS Humanized (Sapiens^3)
VQLVESGGGLVQPGGSLRLSCAASGFTVSSNYMSWVRQAPGKGLEWVSVIYSGGSTYYADSVKFTISRDNSKNTLYLQMNSLRAEDTAVYYCAR------AEYFQHWGQGTLVTVSS IGHV3-66*01, IGHJ1*01
```

#### Light chain

```
DIQMTQSPSSLSASVGDRVTITCRASQSISSYLNWYQQKPGKAPKLLIYAASSLQSGVPSRFSGSGSGTDFTLTISSLQPEDFATYYCQQSYST----LTFGGGTKVEIK IGKV1-39*01, IGKJ4*01
AIQMTQSPSSLSASVGDRVTITCQASQSINNELSWYQQKPGKAPKLLIYRASTLASGVPSRFSGSGSGTDFTLTISSLQPDDFATYYCQQGYSLRNIDNAFGGGTKVEIK Humanized (Experimental)
AYDMTQTPASVSAAVGGTVTIKCQASQSINNELSWYQQKPGQRPKLLIYRASTLASGVSSRFKGSGSGTEFTLTISDLECADAATYYCQQGYSLRNIDNAFGGGTEVVVK Parental
DIQMTQSPSSLSASVGDRVTITCQASQSINNELSWYQQKPGKAPKLLIYRASTLASGVPSRFSGSGSGTEFTLTISSLQPEDFATYYCQQGYSLRNIDNAFGGGTKVEIK Humanized (Sapiens^3)
DIQMTQSPSSLSASVGDRVTITCRASQSISSYLNWYQQKPGKAPKLLIYAASSLQSGVPSRFSGSGSGTDFTLTISSLQPEDFATYYCQQSYST----LTFGGGTKVEIK IGKV1-39*01, IGKJ4*01
```

### Crizanlizumab

#### Heavy chain

```
QVQLVQSGAEVKKPGASVKVSCKASGYTFTSYYMHWVRQAPGQGLEWMGIINPSGGSTSYAQKFQGRVTMTRDTSTSTVYMELSSLRSEDTAVYYCAR---------YFDYWGQGTLVTVSS IGHV1-46*01, IGHJ4*01
QVQLVQSGAEVKKPGASVKVSCKVSGYTFTSYDINWVRQAPGKGLEWMGWIYPGDGSIKYNEKFKGRVTMTVDKSTDTAYMELSSLRSEDTAVYYCARRGEYGNYEGAMDYWGQGTLVTVSS Humanized (Experimental)
QVQLQQSGPELVKPGALVKISCKASGYTFTSYDINWVKQRPGQGLEWIGWIYPGDGSIKYNEKFKGKATLTVDKSSSTAYMQVSSLTSENSAVYFCARRGEYGNYEGAMDYWGQGTTVTVSS Parental
QVQLVQSGPEVKKPGASVKISCKASGYTFTSYDINWVRQAPGQGLEWMGWIYPGDGSIKYNEKFKGRVTLTRDTSASTAYMEVSSLTSEDTAVYFCARRGEYGNYEGAMDYWGQGTTVTVSS Humanized (Sapiens^3)
QVQLVQSGAEVKKPGASVKVSCKASGYTFTSYAMHWVRQAPGQRLEWMGWINAGNGNTKYSQKFQGRVTITRDTSASTAYMELSSLRSEDTAVYYCAR--------DAFDVWGQGTMVTVSS IGHV1-3*01, IGHJ3*01
```

#### Light chain

```
DIQMTQSPSSLSASVGDRVTITCRASQSI----SSYLNWYQQKPGKAPKLLIYAASSLQSGVPSRFSGSGSGTDFTLTISSLQPEDFATYYCQQSYS--LTFGGGTKVEIK IGKV1-39*01, IGKJ4*01
DIQMTQSPSSLSASVGDRVTITCKASQSVDYDGHSYMNWYQQKPGKAPKLLIYAASNLESGVPSRFSGSGSGTDFTLTISSLQPEDFATYYCQQSDENPLTFGGGTKVEIK Humanized (Experimental)
DIVLTQSPASLAVSLGQRATISCKASQSVDYDGHSYMNWYQQKPGQPPKLLIYAASNLESGIPARFSGSGSGTDFTLNIHPVEEEDAATYYCQQSDENPLTFGTGTKLELK Parental
DIVLTQSPASLAVSPGERATISCKASQSVDYDGHSYMNWYQQKPGQPPKLLIYAASNLESGVPDRFSGSGSGTDFTLTISRVEAEDVAVYYCQQSDENPLTFGQGTKLEIK Humanized (Sapiens^3)
DIVMTQSPDSLAVSLGERATINCKSSQSVLYNNKNYLAWYQQKPGQPPKLLIYWASTRESGVPDRFSGSGSGTDFTLTISSLQAEDVAVYYCQQYYS--YTFGQGTKLEIK IGKV4-1*01, IGKJ2*01
```

### Eculizumab

#### Heavy chain

```
QVQLVQSGAEVKKPGASVKVSCKASGYTFTSYYMHWVRQAPGQGLEWMGIINPSGGSTSYAQKFQGRVTMTRDTSTSTVYMELSSLRSEDTAVYYCAR-------YWYFDLWGRGTLVTVSS IGHV1-46*01, IGHJ2*01
QVQLVQSGAEVKKPGASVKVSCKASGYIFSNYWIQWVRQAPGQGLEWMGEILPGSGSTEYTENFKDRVTMTRDTSTSTVYMELSSLRSEDTAVYYCARYFFGSSPNWYFDVWGQGTLVTVSS Humanized (Experimental)
QVQLQQSGAELMKPGASVKMSCKATGYIFSNYWIQWIKQRPGHGLEWIGEILPGSGSTEYTENFKDKAAFTADTSSNTAYMQLSSLTSEDSAVYYCARYFFGSSPNWYFDVWGAGTTVTVSS Parental
QVQLVQSGAEVKKPGASVKISCKASGYIFSNYWIQWVRQAPGQGLEWMGEILPGSGSTEYTENFKDRVTFTADTSTNTAYMELSSLTSEDTAVYYCARYFFGSSPNWYFDVWGRGTLVTVSS Humanized (Sapiens^3)
QVQLVQSGAEVKKPGASVKVSCKASGYTFTSYYMHWVRQAPGQGLEWMGIINPSGGSTSYAQKFQGRVTMTRDTSTSTVYMELSSLRSEDTAVYYCAR-------YWYFDLWGRGTLVTVSS IGHV1-46*01, IGHJ2*01
```

#### Light chain

```
DIQMTQSPSSLSASVGDRVTITCQASQDISNYLNWYQQKPGKAPKLLIYDASNLETGVPSRFSGSGSGTDFTFTISSLQPEDIATYYCQQYDN--WTFGQGTKVEIK IGKV1-33*01, IGKJ1*01
DIQMTQSPSSLSASVGDRVTITCGASENIYGALNWYQQKPGKAPKLLIYGATNLADGVPSRFSGSGSGTDFTLTISSLQPEDFATYYCQNVLNTPLTFGQGTKVEIK Humanized (Experimental)
DIQMTQSPASLSASVGETVTITCGASENIYGALNWYQRKQGKSPQLLIYGATNLADGMSSRFSGSGSGRQYYLKISSLHPDDVATYYCQNVLNTPLTFGAGTKLELK Parental
DIQMTQSPSSLSASVGDRVTITCGASENIYGALNWYQQKPGKAPKLLIYGATNLADGVPSRFSGSGSGTDFTLTISSLQPEDVATYYCQNVLNTPLTFGQGTKLEIK Humanized (Sapiens^3)
DIQMTQSPSSLSASVGDRVTITCQASQDISNYLNWYQQKPGKAPKLLIYDASNLETGVPSRFSGSGSGTDFTFTISSLQPEDIATYYCQQYDN--YTFGQGTKLEIK IGKV1-33*01, IGKJ2*01
```

### Etaracizumab

#### Heavy chain

```
QVQLVESGGGVVQPGRSLRLSCAASGFTFSSYAMHWVRQAPGKGLEWVAVISYDGSNKYYADSVKGRFTISRDNSKNTLYLQMNSLRAEDTAVYYCAR---NWFDSWGQGTLVTVSS IGHV3-30*01, IGHJ5*01
QVQLVESGGGVVQPGRSLRLSCAASGFTFSSYDMSWVRQAPGKGLEWVAKVSSGGGSTYYLDTVQGRFTISRDNSKNTLYLQMNSLRAEDTAVYYCARHLHGSFASWGQGTTVTVSS Humanized (Experimental)
EVQLEESGGGLVKPGGSLKLSCAASGFAFSSYDMSWVRQIPEKRLEWVAKVSSGGGSTYYLDTVQGRFTISRDNAKNTLYLQMSSLNSEDTAMYYCARHNYGSFAYWGQGTLVTVSA Parental
EVQLVESGGGLVKPGGSLRLSCAASGFTFSSYDMSWVRQAPGKGLEWVSKVSSGGGSTYYLDTVQGRFTISRDNAKNSLYLQMNSLRAEDTAVYYCARHNYGSFAYWGQGTLVTVSS Humanized (Sapiens^3)
EVQLVESGGGLVKPGGSLRLSCAASGFTFSSYSMNWVRQAPGKGLEWVSSISSSSSYIYYADSVKGRFTISRDNAKNSLYLQMNSLRAEDTAVYYCAR----YFDYWGQGTLVTVSS IGHV3-21*01, IGHJ4*01
```

#### Light chain

```
EIVLTQSPATLSLSPGERATLSCRASQSVSSYLAWYQQKPGQAPRLLIYDASNRATGIPARFSGSGSGTDFTLTISSLEPEDFAVYYCQQRSN--LTFGGGTKVEIK IGKV3-11*01, IGKJ4*01
EIVLTQSPATLSLSPGERATLSCQASQSISNFLHWYQQRPGQAPRLLIRYRSQSISGIPARFSGSGSGTDFTLTISSLEPEDFAVYYCQQSGSWPLTFGGGTKVEIK Humanized (Experimental)
ELVMTQTPATLSVTPGDSVSLSCRASQSISNHLHWYQQKSHESPRLLIKYASQSISGIPSRFSGSGSGTDFTLSINSVETEDFGMYFCQQSNSWPHTFGGGTKLEIK Parental
EIVMTQSPATLSVSPGERVTLSCRASQSISNHLHWYQQKPGQAPRLLIYYASQSISGIPARFSGSGSGTDFTLTISSLEPEDFAVYYCQQSNSWPHTFGQGTKLEIK Humanized (Sapiens^3)
EIVLTQSPATLSLSPGERATLSCRASQSVSSYLAWYQQKPGQAPRLLIYDASNRATGIPARFSGSGSGTDFTLTISSLEPEDFAVYYCQQRSN--YTFGQGTKLEIK IGKV3-11*01, IGKJ2*01
```

### Herceptin

#### Heavy chain

```
EVQLVESGGGLVQPGGSLRLSCAASGFTVSSNYMSWVRQAPGKGLEWVSVIYSG-GSTYYADSVKGRFTISRDNSKNTLYLQMNSLRAEDTAVYYCAR-------YFDYWGQGTLVTVSS IGHV3-66*01, IGHJ4*01
EVQLVESGGGLVQPGGSLRLSCAASGFNIKDTYIHWVRQAPGKGLEWVARIYPTNGYTRYADSVKGRFTISADTSKNTAYLQMNSLRAEDTAVYYCSRWGGDGFYAMDYWGQGTLVTVSS Humanized (Experimental)
QVQLQQSGPELVKPGASLKLSCTASGFNIKDTYIHWVKQRPEQGLEWIGRIYPTNGYTRYDPKFQDKATITADTSSNTAYLQVSRLTSEDTAVYYCSRWGGDGFYAMDYWGQGASVTVSS Parental
QVQLVQSGPEVKKPGASVKVSCKASGYTFTDTYIHWVRQAPGQGLEWMGRIYPTNGYTRYDPKFQDRVTITRDTSINTAYLQVSRLTSDDTAVYYCARWGGDGFYAMDYWGQGTLVTVSS Humanized (Sapiens^3)
QVQLVQSGAEVKKPGASVKVSCKASGYTFTGYYMHWVRQAPGQGLEWMGRINPNSGGTNYAQKFQGRVTMTRDTSISTAYMELSRLRSDDTAVYYCAR-------YFDYWGQGTLVTVSS IGHV1-2*06, IGHJ4*01
```

#### Light chain

```
AIQLTQSPSSLSASVGDRVTITCRASQGISSALAWYQQKPGKAPKLLIYDASSLESGVPSRFSGSGSGTDFTLTISSLQPEDFATYYCQQFNS--WTFGQGTKVEIK IGKV1-13*02, IGKJ1*01
DIQMTQSPSSLSASVGDRVTITCRASQDVNTAVAWYQQKPGKAPKLLIYSASFLESGVPSRFSGSRSGTDFTLTISSLQPEDFATYYCQQHYTTPTTFGQGTKVEIK Humanized (Experimental)
DIVMTQSHKFMSTSVGDRVSITCKASQDVNTAVAWYQQKPGHSPKLLIYSASFRYTGVPDRFTGNRSGTDFTFTISSVQAEDLAVYYCQQHYTTPPTFGGGTKVEIK Parental
DIQMTQSPSFLSASVGDRVTITCKASQDVNTAVAWYQQKPGKAPKLLIYSASFRYTGVPSRFSGSGSGTDFTLTISSLQPEDFATYYCQQHYTTPPTFGGGTKVEIK Humanized (Sapiens^3)
DIQMTQSPSSLSASVGDRVTITCRASQSISSYLNWYQQKPGKAPKLLIYAASSLQSGVPSRFSGSGSGTDFTLTISSLQPEDFATYYCQQSYS--LTFGGGTKVEIK IGKV1-39*01, IGKJ4*01
```

### Idarucizumab

#### Heavy chain

```
QVQLQESGPGLVKPSETLSLTCTVSGGSISSYYWSWIRQPPGKGLEWIGYIYTSGSTNYNPSLKSRVTISVDTSKNQFSLKLSSVTAADTAVYYCAR----------YFDYWGQGTLVTVSS IGHV4-4*08, IGHJ4*01
QVQLQESGPGLVKPSETLSLTCTVSGFSLTSYIVDWIRQPPGKGLEWIGVIWAGGSTGYNSALRSRVSITKDTSKNQFSLKLSSVTAADTAVYYCASAAYYSYYNYDGFAYWGQGTLVTVSS Humanized (Experimental)
QVQLEQSGPGLVAPSQRLSITCTVSGFSLTSYIVDWVRQSPGKGLEWLGVIWAGGSTGYNSALRSRLSITKSNSKSQVFLQMNSLQTDDTAIYYCASAAYYSYYNYDGFAYWGQGTLVTVSA Parental
QVQLQESGPGLVKPSETLSLTCTVSGGSLTSYIVDWVRQSPGKGLEWLGVIWAGGSTGYNSALRSRLSISLDTSKSQVFLQLNSLTAADTAIYYCARAAYYSYYNYDGFAYWGQGTLVTVSS Humanized (Sapiens^3)
QVQLQESGPGLVKPSETLSLTCTVSGGSISSYYWSWIRQPPGKGLEWIGYIYTSGSTNYNPSLKSRVTISVDTSKNQFSLKLSSVTAADTAVYYCAR----------YFDYWGQGTLVTVSS IGHV4-4*08, IGHJ4*01
```

#### Light chain

```
DVVMTQSPLSLPVTLGQPASISCRSSQSLVYSDGNTYLNWFQQRPGQSPRRLIYKVSNRDSGVPDRFSGSGSGTDFTLKISRVEAEDVGVYYCMQGTH--LTFGGGTKVEIK IGKV2-30*01, IGKJ4*01
DVVMTQSPLSLPVTLGQPASISCKSSQSLLYTDGKTYLYWFLQRPGQSPRRLIYLVSKLDSGVPDRFSGSGSGTDFTLKISRVEAEDVGVYYCLQSTHFPHTFGGGTKVEIK Humanized (Experimental)
DVVMTQTPLTLSVTIGQPASISCKSSQSLLYTNGKTYLYWLLQRPGQSPKRLIYLVSKLDSGVPDRFSGSGSGTDFTLKISRVEAEDVGIYYCLQSTHFPHTFGGGTKLEIK Parental
DVVMTQSPLSLSVTPGQPASISCKSSQSLLYTNGKTYLYWFQQRPGQSPRRLIYLVSKLDSGVPDRFSGSGSGTDFTLKISRVEAEDVGIYYCLQSTHFPHTFGQGTKLEIK Humanized (Sapiens^3)
DVVMTQSPLSLPVTLGQPASISCRSSQSLVYSDGNTYLNWFQQRPGQSPRRLIYKVSNRDSGVPDRFSGSGSGTDFTLKISRVEAEDVGVYYCMQGTH--YTFGQGTKLEIK IGKV2-30*01, IGKJ2*01
```

### Ixekizumab

#### Heavy chain

```
QVQLVQSGAEVKKPGSSVKVSCKASGGTFSSYAISWVRQAPGQGLEWMGGIIPIFGTANYAQKFQGRVTITADESTSTAYMELSSLRSEDTAVYYCAR------YFDYWGQGTLVTVSS IGHV1-69*01, IGHJ4*01
QVQLVQSGAEVKKPGSSVKVSCKASGYSFTDYHIHWVRQAPGQGLEWMGVINPMYGTTDYNQRFKGRVTITADESTSTAYMELSSLRSEDTAVYYCARYDYFTGTGVYWGQGTLVTVSS Humanized (Experimental)
QVQLQQSRPELVKPGASVKISCKASGYSFTDYNMNWVKQSNGKSLEWIGVINPNYGTTDYNQRFKGKATLTVDQSSRTAYMQLNSLTSEDSAVYYCVIYDYATGTGGYWGQGSPLTVSS Parental
QVQLVQSGPELKKPGASVKVSCKASGYSFTDYNMNWVRQAPGQGLEWMGVINPNYGTTDYNQRFKGRVTLTRDTSIRTAYMELNSLTSEDTAVYYCVIYDYATGTGGYWGQGTLVTVSS Humanized (Sapiens^3)
QVQLVQSGAEVKKPGASVKVSCKASGYTFTGYYMHWVRQAPGQGLEWMGWINPNSGGTNYAQKFQGRVTMTRDTSISTAYMELSRLRSDDTAVYYCAR------YFDYWGQGTLVTVSS IGHV1-2*02, IGHJ4*01
```

#### Light chain

```
DIVMTQTPLSLSVTPGQPASISCKSSQSLLHSDGKTYLYWYLQKPGQSPQLLIYEVSNRFSGVPDRFSGSGSGTDFTLKISRVEAEDVGVYYCMQSIQ--YTFGQGTKLEIK IGKV2D-29*02, IGKJ2*01
DIVMTQTPLSLSVTPGQPASISCRSSRSLVHSRGNTYLHWYLQKPGQSPQLLIYKVSNRFIGVPDRFSGSGSGTDFTLKISRVEAEDVGVYYCSQSTHLPFTFGQGTKLEIK Humanized (Experimental)
DVVLTQTPLSLPVSLGDQASISCRSSQSLVHSNGNTYLHWYLQKPGQSPKLLIYKVSNRFSGVPDRFSGSGSGTDFTLKISRVEAEDLGVYFCSQSTHVPFTFGSGTKLEIK Parental
DIVMTQTPLSLPVTLGQPASISCRSSQSLVHSNGNTYLHWYLQKPGQSPQLLIYKVSNRFSGVPDRFSGSGSGTDFTLKISRVEAEDVGVYYCSQSTHVPFTFGQGTKLEIK Humanized (Sapiens^3)
DIVMTQTPLSLSVTPGQPASISCKSSQSLLHSDGKTYLYWYLQKPGQSPQLLIYEVSNRFSGVPDRFSGSGSGTDFTLKISRVEAEDVGVYYCMQSIQ--YTFGQGTKLEIK IGKV2D-29*02, IGKJ2*01
```

### Ligelizumab

#### Heavy chain

```
QVQLVQSGAEVKKPGSSVKVSCKASGGTFSSYAISWVRQAPGQGLEWMGGIIPIFGTANYAQKFQGRVTITADESTSTAYMELSSLRSEDTAVYYCAR----------YFDYWGQGTLVTVSS IGHV1-69*01, IGHJ4*01
QVQLVQSGAEVMKPGSSVKVSCKASGYTFSWYWLEWVRQAPGHGLEWMGEIDPGTFTTNYNEKFKARVTFTADTSTSTAYMELSSLRSEDTAVYYCARFSHFSGSNYDYFDYWGQGTLVTVSS Humanized (Experimental)
QVQLQQSGAELMKPGASVKISCKTTGYTFSMYWLEWVKQRPGHGLEWVGEISPGTFTTNYNEKFKAKATFTADTSSNTAYLQLSGLTSEDSAVYFCARFSHFSGSNYDYFDYWGQGTSLTVSS Parental
QVQLVQSGAEVKKPGASVKISCKTSGYTFSMYWLEWVRQAPGQGLEWVGEISPGTFTTNYNEKFKARVTFTADTSTNTAYLELSGLTSEDTAVYFCARFSHFSGSNYDYFDYWGQGTLVTVSS Humanized (Sapiens^3)
QVQLVQSGAEVKKPGSSVKVSCKASGGTFSSYAISWVRQAPGQGLEWMGGIIPIFGTANYAQKFQGRVTITADESTSTAYMELSSLRSEDTAVYYCAR----------YFDYWGQGTLVTVSS IGHV1-69*01, IGHJ4*01
```

#### Light chain

```
EIVMTQSPATLSVSPGERATLSCRASQSVSSNLAWYQQKPGQAPRLLIYGASTRATGIPARFSGSGSGTEFTLTISSLQSEDFAVYYCQQYNN--LTFGGGTKVEIK IGKV3-15*01, IGKJ4*01
EIVMTQSPATLSVSPGERATLSCRASQSIGTNIHWYQQKPGQAPRLLIYYASESISGIPARFSGSGSGTEFTLTISSLQSEDFAVYYCQQSWSWPTTFGGGTKVEIK Humanized (Experimental)
DILLTQSPAILSVSPGERVSFSCRASQSIGTNIHWYQQRTDGSPRLLIKYASESISGIPSRFSGSGSGTEFTLNINSVESEDIADYYCQQSDSWPTTFGGGTKLEIK Parental
DIVLTQSPATLSVSPGERVTLSCRASQSIGTNIHWYQQRPGQAPRLLIYYASESISGIPARFSGSGSGTEFTLTINSLQSEDFAVYYCQQSDSWPTTFGQGTKVEIK Humanized (Sapiens^3)
EIVMTQSPATLSVSPGERATLSCRASQSVSSNLAWYQQKPGQAPRLLIYGASTRATGIPARFSGSGSGTEFTLTISSLQSEDFAVYYCQQYNN--WTFGQGTKVEIK IGKV3-15*01, IGKJ1*01
```

### Lorvotuzumab

#### Heavy chain

```
QVQLVESGGGVVQPGRSLRLSCAASGFTFSSYGMHWVRQAPGKGLEWVAVISYDGSNKYYADSVKGRFTISRDNSKNTLYLQMNSLRAEDTAVYYCAR-----YFDYWGQGTLVTVSS IGHV3-30*03, IGHJ4*01
QVQLVESGGGVVQPGRSLRLSCAASGFTFSSFGMHWVRQAPGKGLEWVAYISSGSFTIYYADSVKGRFTISRDNSKNTLYLQMNSLRAEDTAVYYCARMRKGYAMDYWGQGTLVTVSS Humanized (Experimental)
DVQLVESGGGLVQPGGSRKLSCAASGFTFSSFGMHWVRQAPEKGLEWVAYISSGSFTIYHADTVKGRFTISRDNPKNTLFLQMTSLRAEDTAHYYCARMRKGYAMDYWGQGTTVTVSS Parental
EVQLVESGGGLVQPGGSLRLSCAASGFTFSSFGMHWVRQAPGKGLEWVSYISSGSFTIYHADTVKGRFTISRDNAKNSLYLQMNSLRAEDTAVYYCARMRKGYAMDYWGQGTTVTVSS Humanized (Sapiens^3)
EVQLVESGGGLVQPGGSLRLSCAASGFTFSSYSMNWVRQAPGKGLEWVSYISSSSSTIYYADSVKGRFTISRDNAKNSLYLQMNSLRAEDTAVYYCAR----DAFDVWGQGTMVTVSS IGHV3-48*01, IGHJ3*01
```

#### Light chain

```
DVVMTQSPLSLPVTLGQPASISCRSSQSLVHSDGNTYLNWFQQRPGQSPRRLIYKVSNRDSGVPDRFSGSGSGTDFTLKISRVEAEDVGVYYCMQGTH--WTFGQGTKVEIK IGKV2-30*02, IGKJ1*01
DVVMTQSPLSLPVTLGQPASISCRSSQIIIHSDGNTYLEWFQQRPGQSPRRLIYKVSNRFSGVPDRFSGSGSGTDFTLKISRVEAEDVGVYYCFQGSHVPHTFGQGTKVEIK Humanized (Experimental)
DVLMTQTPLSLPVSLGDQASISCRSSQIIIHSDGNTYLEWFLQKPGQSPKLLIYKVSNRFSGVPDRFSGSGSGTDFTLMISRVEAEDLGVYYCFQGSHVPHTFGGGTKLEIK Parental
DVVMTQTPLSLPVTLGQPASISCRSSQIIIHSDGNTYLEWYLQKPGQSPQLLIYKVSNRFSGVPDRFSGSGSGTDFTLKISRVEAEDVGVYYCFQGSHVPHTFGQGTKLEIK Humanized (Sapiens^3)
DIVMTQTPLSLSVTPGQPASISCKSSQSLLHSDGKTYLYWYLQKPGQSPQLLIYEVSSRFSGVPDRFSGSGSGTDFTLKISRVEAEDVGVYYCMQGIH--YTFGQGTKLEIK IGKV2-29*02, IGKJ2*01
```

### Mogamulizumab

#### Heavy chain

```
EVQLVESGGGLVKPGGSLRLSCAASGFTFSSYSMNWVRQAPGKGLEWVSSISSSSSYIYYADSVKGRFTISRDNAKNSLYLQMNSLRAEDTAVYYCAR------YFDYWGQGTLVTVSS IGHV3-21*01, IGHJ4*01
EVQLVESGGDLVQPGRSLRLSCAASGFIFSNYGMSWVRQAPGKGLEWVATISSASTYSYYPDSVKGRFTISRDNAKNSLYLQMNSLRVEDTALYYCGRHSDGNFAFGYWGQGTLVTVSS Humanized (Experimental)
EVQLVESGGDLMKPGGSLKISCAASGFIFSNYGMSWVRQTPDMRLEWVATISSASTYSYYPDSVKGRFTISRDNAENSLYLQMNSLRSEDTGIYYCGRHSDGNFAFGYWGRGTLVTVSA Parental
EVQLVESGGGLVKPGGSLRLSCAASGFTFSNYGMSWVRQAPGKGLEWVSTISSASTYSYYPDSVKGRFTISRDNAKNSLYLQMNSLRAEDTAVYYCARHSDGNFAFGYWGQGTLVTVSS Humanized (Sapiens^3)
EVQLVESGGGLVKPGGSLRLSCAASGFTFSSYSMNWVRQAPGKGLEWVSSISSSSSYIYYADSVKGRFTISRDNAKNSLYLQMNSLRAEDTAVYYCAR------YFDYWGQGTLVTVSS IGHV3-21*01, IGHJ4*01
```

#### Light chain

```
DIVMTQSPLSLPVTPGEPASISCRSSQSLLHSNGYNYLDWYLQKPGQSPQLLIYLGSNRASGVPDRFSGSGSGTDFTLKISRVEAEDVGVYYCMQALQ--WTFGQGTKVEIK IGKV2-28*01, IGKJ1*01
DVLMTQSPLSLPVTPGEPASISCRSSRNIVHINGDTYLEWYLQKPGQSPQLLIYKVSNRFSGVPDRFSGSGSGTDFTLKISRVEAEDVGVYYCFQGSLLPWTFGQGTKVEIK Humanized (Experimental)
DVLMTQTPLSLPVSLGDQASISCRSSRNIVHINGDTYLEWYLQRPGQSPKLLIYKVSNRFSGVPDRFSGSGSGTDFTLKISRVEAEDLGVYYCFQGSLLPWTFGGGTRLEIR Parental
DVVMTQSPLSLPVTLGQPASISCRSSRNIVHINGDTYLEWYLQRPGQSPQLLIYKVSNRFSGVPDRFSGSGSGTDFTLKISRVEAEDVGVYYCFQGSLLPWTFGQGTRLEIK Humanized (Sapiens^3)
DVVMTQSPLSLPVTLGQPASISCRSSQSLVHSDGNTYLNWFQQRPGQSPRRLIYKVSNRDSGVPDRFSGSGSGTDFTLKISRVEAEDVGVYYCMQGTH--ITFGQGTRLEIK IGKV2-30*02, IGKJ5*01
```

### Omalizumab

#### Heavy chain

```
EVQLVESGGGLVQPGGSLRLSCAASGFTV-SSNYMSWVRQAPGKGLEWVSVIYSGGSTYYADSVKGRFTISRDNSKNTLYLQMNSLRAEDTAVYYCAR------AEYFQHWGQGTLVTVSS IGHV3-66*01, IGHJ1*01
EVQLVESGGGLVQPGGSLRLSCAVSGYSITSGYSWNWIRQAPGKGLEWVASITYDGSTNYADSVKGRFTISRDDSKNTFYLQMNSLRAEDTAVYYCARGSHYFGHWHFAVWGQGTLVTVSS Humanized (Experimental)
DVQLQESGPGLVKPSQSLSLACSVTGYSITSGYSWNWIRQFPGNKLEWMGSITYDGSSNYNPSLKNRISVTRDTSQNQFFLKLNSATAEDTATYYCARGSHYFGHWHFAVWGAGTTVTVSS Parental
QVQLQESGPGLVKPSQTLSLTCSVSGYSITSGYSWNWIRQPPGKGLEWIGSITYDGSSNYNPSLKNRVSVSVDTSKNQFSLKLNSVTAADTAVYYCARGSHYFGHWHFAVWGKGTTVTVSS Humanized (Sapiens^3)
QVQLQESGPGLVKPSETLSLTCAVSGYSISSGYYWGWIRQPPGKGLEWIGSIYHSGSTYYNPSLKSRVTISVDTSKNQFSLKLSSVTAADTAVYYCAR------YWYFDLWGRGTLVTVSS IGHV4-38-2*01, IGHJ2*01
```

#### Light chain

```
DIQMTQSPSSLSASVGDRVTITCRASQSI----SSYLNWYQQKPGKAPKLLIYAASSLQSGVPSRFSGSGSGTDFTLTISSLQPEDFATYYCQQSYS--WTFGQGTKVEIK IGKV1-39*01, IGKJ1*01
DIQLTQSPSSLSASVGDRVTITCRASQSVDYDGDSYMNWYQQKPGKAPKLLIYAASYLESGVPSRFSGSGSGTDFTLTISSLQPEDFATYYCQQSHEDPYTFGQGTKVEIK Humanized (Experimental)
DIQLTQSPASLAVSLGQRATISCKASQSVDYDGDSYMNWYQQKPGQPPILLIYAASYLGSEIPARFSGSGSGTDFTLNIHPVEEEDAATFYCQQSHEDPYTFGAGTKLEIK Parental
DIVLTQSPDSLAVSPGERATISCKASQSVDYDGDSYMNWYQQKPGQPPKLLIYAASYLGSGVPDRFSGSGSGTDFTLTISRVEAEDVAVYYCQQSHEDPYTFGQGTKLEIK Humanized (Sapiens^3)
DIVMTQSPDSLAVSLGERATINCKSSQSVLYNNKNYLAWYQQKPGQPPKLLIYWASTRESGVPDRFSGSGSGTDFTLTISSLQAEDVAVYYCQQYYS--YTFGQGTKLEIK IGKV4-1*01, IGKJ2*01
```

### Palivizumab

#### Heavy chain

```
QVTLRESGPALVKPTQTLTLTCTFSGFSLSTSGMCVSWIRQPPGKALEWLALIDWDDDKYYSTSLKTRLTISKDTSKNQVVLTMTNMDPVDTATYYCARI---YWYFDLWGRGTLVTVSS IGHV2-70*01, IGHJ2*01
QVTLRESGPALVKPTQTLTLTCTFSGFSLSTSGMSVGWIRQPPGKALEWLADIWWDDKKDYNPSLKSRLTISKDTSKNQVVLKVTNMDPADTATYYCARSMITNWYFDVWGAGTTVTVSS Humanized (Experimental)
QVELQESGPGILQPSQTLSLTCSFSGFSLSTSGMSVGWIRQPSGEGLEWLADIWWDDKKDYNPSLKSRLTISKDTSSNQVFLKITGVDTADTATYYCARSMITNWYFDVWGAGTTVTVSS Parental
QVTLKESGPTLVKPTQTLTLTCTFSGFSLSTSGMSVGWIRQPPGKALEWLADIWWDDKKDYNPSLKSRLTITKDTSKNQVVLTMTNMDPVDTATYYCARSMITNWYFDVWGRGTLVTVSS Humanized (Sapiens^3)
QVTLKESGPTLVKPTQTLTLTCTFSGFSLSTSGVGVGWIRQPPGKALEWLALIYWDDDKRYGPSLKSRLTITKDTSKNQVVLTMTNMDPVDTATYYCAHR---YWYFDLWGRGTLVTVSS IGHV2-5*09, IGHJ2*01
```

#### Light chain

```
DIQMTQSPSTLSASVGDRVTITCRASQSISWLAWYQQKPGKAPKLLIYDASSLESGVPSRFSGSGSGTEFTLTISSLQPDDFATYYCQQYNS--YTFGQGTKLEIK IGKV1-5*01, IGKJ2*01
DIQMTQSPSTLSASVGDRVTITCKCQLSVGYMHWYQQKPGKAPKLLIYDTSKLASGVPSRFSGSGSGTEFTLTISSLQPDDFATYYCFQGSGYPFTFGGGTKLEIK Humanized (Experimental)
DIQLTQSPAIMSASPGEKVTMTCSASSSVGYMHWYQQKSSTSPKLWIYDTSKLASGVPGRFSGSGSGNSYSLTISSIQAEDVATYYCFQGSGYPFTFGQGTKLEIK Parental
DIVLTQSPATLSLSPGERATLSCSASSSVGYMHWYQQKPGKAPKLLIYDTSKLASGVPSRFSGSGSGTDFTLTISSLQAEDVATYYCFQGSGYPFTFGQGTKLEIK Humanized (Sapiens^3)
EIVLTQSPATLSLSPGERATLSCRASQSVSYLAWYQQKPGQAPRLLIYDASNRATGIPARFSGSGSGTDFTLTISSLEPEDFAVYYCQQRSN--YTFGQGTKLEIK IGKV3-11*01, IGKJ2*01
```

### Pembrolizumab

#### Heavy chain

```
QVQLVQSGAEVKKPGASVKVSCKASGYTFTGYYMHWVRQAPGQGLEWMGWINPNSGGTNYAQKFQGRVTMTRDTSISTAYMELSRLRSDDTAVYYCAR-------YFDYWGQGTLVTVSS IGHV1-2*02, IGHJ4*01
QVQLVQSGVEVKKPGASVKVSCKASGYTFTNYYMYWVRQAPGQGLEWMGGINPSNGGTNFNEKFKNRVTLTTDSSTTTAYMELKSLQFDDTAVYYCARRDYRFDMGFDYWGQGTTVTVSS Humanized (Experimental)
QVQLQQPGAELVKPGTSVKLSCKASGYTFTNYYMYWVKQRPGQGLEWIGGINPSNGGTNFNEKFKNKATLTVDSSSSTTYMQLSSLTSEDSAVYYCTRRDYRFDMGFDYWGQGTTLTVSS Parental
QVQLVQSGAEVKKPGASVKVSCKASGYTFTNYYMYWVRQAPGQGLEWMGGINPSNGGTNFNEKFKNRVTLTRDTSTSTAYMELSSLTSEDTAVYYCARRDYRFDMGFDYWGQGTLVTVSS Humanized (Sapiens^3)
QVQLVQSGAEVKKPGASVKVSCKASGYTFTSYYMHWVRQAPGQGLEWMGIINPSGGSTSYAQKFQGRVTMTRDTSTSTVYMELSSLRSEDTAVYYCAR-------YFDYWGQGTLVTVSS IGHV1-46*01, IGHJ4*01
```

#### Light chain

```
EIVLTQSPATLSLSPGERATLSCRASQSV----SSYLAWYQQKPGQAPRLLIYDASNRATGIPARFSGSGSGTDFTLTISSLEPEDFAVYYCQQRSN--LTFGGGTKVEIK IGKV3-11*01, IGKJ4*01
EIVLTQSPATLSLSPGERATLSCRASKGVSTSGYSYLHWYQQKPGQAPRLLIYLASYLESGVPARFSGSGSGTDFTLTISSLEPEDFAVYYCQHSRDLPLTFGGGTKVEIK Humanized (Experimental)
DIVLTQSPASLAVSLGQRAAISCRASKGVSTSGYSYLHWYQQKPGQSPKLLIYLASYLESGVPARFSGSGSGTDFTLNIHPVEEEDAATYYCQHSRDLPLTFGTGTKLELK Parental
DIVLTQSPASLAVSPGERAAISCRASKGVSTSGYSYLHWYQQKPGQSPKLLIYLASYLESGVPDRFSGSGSGTDFTLTISRVEAEDVAVYYCQHSRDLPLTFGQGTKLEIK Humanized (Sapiens^3)
DIVLTQSPASLAVSPGQRATITCRASESVSFLGINLIHWYQQKPGQPPKLLIYQASNKDTGVPARFSGSGSGTDFTLTINPVEANDTANYYCLQSKN--YTFGQGTKLEIK IGKV7-3*01, IGKJ2*01
```

### Pertuzumab

#### Heavy chain

```
EVQLVESGGGLVQPGGSLRLSCAASGFTFSSYAMSWVRQAPGKGLEWVSAISGSGGSTYYADSVKGRFTISRDNSKNTLYLQMNSLRAEDTAVYYCAK------YFDYWGQGTLVTVSS IGHV3-23*04, IGHJ4*01
EVQLVESGGGLVQPGGSLRLSCAASGFTFTDYTMDWVRQAPGKGLEWVADVNPNSGGSIYNQRFKGRFTLSVDRSKNTLYLQMNSLRAEDTAVYYCARNLGPSFYFDYWGQGTLVTVSS Humanized (Experimental)
EVQLQQSGPELVKPGTSVKISCKASGFTFTDYTMDWVKQSHGKSLEWIGDVNPNSGGSIYNQRFKGKASLTVDRSSRIVYMELRSLTFEDTAVYYCARNLGPSFYFDYWGQGTTLTVSS Parental
QVQLVQSGPEVKKPGASVKVSCKASGYTFTDYTMDWVRQAPGQGLEWMGDVNPNSGGSIYNQRFKGRVSLTRDTSIRTVYMELRSLTFEDTAVYYCARNLGPSFYFDYWGQGTLVTVSS Humanized (Sapiens^3)
QVQLVQSGAEVKKPGASVKVSCKASGYTFTGYYMHWVRQAPGQGLEWMGWINPNSGGTNYAQKFQGRVTMTRDTSISTAYMELSRLRSDDTAVYYCAR------YFDYWGQGTLVTVSS IGHV1-2*02, IGHJ4*01
```

#### Light chain

```
DIQMTQSPSSLSASVGDRVTITCQASQDISNYLNWYQQKPGKAPKLLIYDASNLETGVPSRFSGSGSGTDFTFTISSLQPEDIATYYCQQYDN--WTFGQGTKVEIK IGKV1-33*01, IGKJ1*01
DIQMTQSPSSLSASVGDRVTITCKASQDVSIGVAWYQQKPGKAPKLLIYSASYRYTGVPSRFSGSGSGTDFTLTISSLQPEDFATYYCQQYYIYPYTFGQGTKVEIK Humanized (Experimental)
DTVMTQSHKIMSTSVGDRVSITCKASQDVSIGVAWYQQRPGQSPKLLIYSASYRYTGVPDRFTGSGSGTDFTFTISSVQAEDLAVYYCQQYYIYPYTFGGGTKLEIK Parental
DIQMTQSPSSLSASVGDRVTITCKASQDVSIGVAWYQQKPGKAPKLLIYSASYRYTGVPSRFSGSGSGTDFTLTISSLQPEDFATYYCQQYYIYPYTFGQGTKLEIK Humanized (Sapiens^3)
DIQMTQSPSSLSASVGDRVTITCQASQDISNYLNWYQQKPGKAPKLLIYDASNLETGVPSRFSGSGSGTDFTFTISSLQPEDIATYYCQQYDN--YTFGQGTKLEIK IGKV1-33*01, IGKJ2*01
```

### Pinatuzumab

#### Heavy chain

```
EVQLVESGGGLVQPGGSLRLSCAASGFTVSSNYMSWVRQAPGKGLEWVSVIYSG-GSTYYADSVKGRFTISRDNSKNTLYLQMNSLRAEDTAVYYCAR-----YWYFDLWGRGTLVTVSS IGHV3-66*01, IGHJ2*01
EVQLVESGGGLVQPGGSLRLSCAASGYEFSRSWMNWVRQAPGKGLEWVGRIYPGDGDTNYSGKFKGRFTISADTSKNTAYLQMNSLRAEDTAVYYCARDGSSWDWYFDVWGQGTLVTVSS Humanized (Experimental)
QVQLQQSGPELVKPGASVKISCKASGYEFSRSWMNWVKQRPGQGREWIGRIYPGDGDTNYSGKFKGKATLTADKSSSTAYMQLSSLTSVDSAVYFCARDGSSWDWYFDVWGAGTTVTVSS Parental
QVQLVQSGPELKKPGASVKISCKASGYTFSRSWMNWVRQAPGQGLEWMGRIYPGDGDTNYSGKFKGRVTLTRDTSTSTAYMELSSLTSEDTAVYFCARDGSSWDWYFDVWGRGTLVTVSS Humanized (Sapiens^3)
QVQLVQSGAEVKKPGASVKVSCKASGYTFTGYYMHWVRQAPGQGLEWMGRINPNSGGTNYAQKFQGRVTMTRDTSISTAYMELSRLRSDDTAVYYCAR-----YWYFDLWGRGTLVTVSS IGHV1-2*06, IGHJ2*01
```

#### Light chain

```
DIQMTQSPSSLSASVGDRVTITCRASQSI-----SSYLNWYQQKPGKAPKLLIYAASSLQSGVPSRFSGSGSGTDFTLTISSLQPEDFATYYCQQSYS--WTFGQGTKVEIK IGKV1-39*01, IGKJ1*01
DIQMTQSPSSLSASVGDRVTITCRSSQSIVHSVGNTFLEWYQQKPGKAPKLLIYKVSNRFSGVPSRFSGSGSGTDFTLTISSLQPEDFATYYCFQGSQFPYTFGQGTKVEIK Humanized (Experimental)
DILMTQTPLSLPVSLGDQASISCRSSQSIVHSNGNTFLEWYLQKPGQSPKLLIYKVSNRFSGVPDRFSGSGSGTDFTLKISRVEAEDLGVYYCFQGSQFPYTFGGGTKVEIK Parental
DIVMTQTPLSLPVTLGQPASISCRSSQSIVHSNGNTFLEWYLQKPGQSPQLLIYKVSNRFSGVPDRFSGSGSGTDFTLKISRVEAEDVGVYYCFQGSQFPYTFGGGTKVEIK Humanized (Sapiens^3)
DIVMTQTPLSLSVTPGQPASISCKSSQSLLHSDGKTYLYWYLQKPGQSPQLLIYEVSNRFSGVPDRFSGSGSGTDFTLKISRVEAEDVGVYYCMQSIQ--LTFGGGTKVEIK IGKV2D-29*02, IGKJ4*01
```

### Refanezumab

#### Heavy chain

```
QVQLVQSGSELKKPGASVKVSCKASGYTFTSYAMNWVRQAPGQGLEWMGWINTNTGNPTYAQGFTGRFVFSLDTSVSTAYLQISSLKAEDTAVYYCAR--------YYYYYGMDVWGQGTTVTVSS IGHV7-4-1*02, IGHJ6*01
QVQLVQSGSELKKPGASVKVSCKASGYTFTNYGMNWVRQAPGQGLEWMGWINTYTGEPTYADDFTGRFVFSLDTSVSTAYLQISSLKAEDTAVYYCARNPINYYGINYEGYVMDYWGQGTLVTVSS Humanized (Experimental)
EIQLVQSGPELKKPGETNKISCKASGYTFTNYGMNWVKQAPGKGLKWMGWINTYTGEPTYADDFTGRFAFSLETSASTAYLQISNLKNEDTATYFCARNPINYYGINYEGYVMDYWGQGTLVTVSS Parental
QVQLVQSGSELKKPGASVKVSCKASGYTFTNYGMNWVRQAPGQGLEWMGWINTYTGEPTYADDFTGRFVFSLDTSVSTAYLQISSLKAEDTAVYYCARNPINYYGINYEGYVMDYWGQGTLVTVSS Humanized (Sapiens^3)
QVQLVQSGSELKKPGASVKVSCKASGYTFTSYAMNWVRQAPGQGLEWMGWINTNTGNPTYAQGFTGRFVFSLDTSVSTAYLQISSLKAEDTAVYYCAR--------YYYYYGMDVWGQGTTVTVSS IGHV7-4-1*02, IGHJ6*01
```

#### Light chain

```
DIVMTQSPDSLAVSLGERATINCKSSQSVLYSSNNKNYLAWYQQKPGQPPKLLIYWASTRESGVPDRFSGSGSGTDFTLTISSLQAEDVAVYYCQQYY--YTFGQGTKLEIK IGKV4-1*01, IGKJ2*01
DIVMTQSPDSLAVSLGERATINCKSSHSVLYSSNQKNYLAWYQQKPGQPPKLLIYWASTRESGVPDRFSGSGSGTDFTLTISSLQAEDVAVYYCHQYLSSLTFGQGTKLEIK Humanized (Experimental)
NIMMTQSPSSLAVSAGEKVTMSCKSSHSVLYSSNQKNYLAWYQQKPGQSPKLLIYWASTRESGVPDRFTGSGSGTDFTLTIINVHTEDLAVYYCHQYLSSLTFGTGTKLEIK Parental
DIVMTQSPDSLAVSLGERATINCKSSHSVLYSSNQKNYLAWYQQKPGQPPKLLIYWASTRESGVPDRFSGSGSGTDFTLTISSLQAEDVAVYYCHQYLSSLTFGQGTKLEIK Humanized (Sapiens^3)
DIVMTQSPDSLAVSLGERATINCKSSQSVLYSSNNKNYLAWYQQKPGQPPKLLIYWASTRESGVPDRFSGSGSGTDFTLTISSLQAEDVAVYYCQQYY--YTFGQGTKLEIK IGKV4-1*01, IGKJ2*01
```

#### Reslizumab

##### Heavy chain

```
EVQLVESGGGLVQPGGSLRLSCAASGFTVSSNYMSWVRQAPGKGLEWVSVIYSGGSTYYADSVKGRFTISRDNSKNTLYLQMNSLRAEDTAVYYCAR----YFDYWGQGTLVTVSS IGHV3-66*01, IGHJ4*01
EVQLVESGGGLVQPGGSLRLSCAVSGLSLTSNSVNWIRQAPGKGLEWVGLIWSNGDTDYNSAIKSRFTISRDTSKSTVYLQMNSLRAEDTAVYYCAREYYGYFDYWGQGTLVTVSS Humanized (Experimental)
EVKLLESGGGLVQPSQTLSLTCTVSGLSLTSNSVNWIRQPPGKGLEWMGLIWSNGDTDYNSAIKSRLSISRDTSKSQVFLKMNSLQSEDTAMYFCAREYYGYFDYWGQGVMVTVSS Parental
QVQLQESGPGLVKPSETLSLTCTVSGGSLTSNSVNWIRQPPGKGLEWIGLIWSNGDTDYNSAIKSRLTISVDTSKSQFFLKLNSVTAADTAMYFCAREYYGYFDYWGQGTLVTVSS Humanized (Sapiens^3)
QVQLQESGPGLVKPSETLSLTCTVSGGSISSYYWSWIRQPPGKGLEWIGYIYTSGSTNYNPSLKSRVTISVDTSKNQFSLKLSSVTAADTAVYYCAR----YFDYWGQGTLVTVSS IGHV4-4*08, IGHJ4*01
```

##### Light chain

```
DIQMTQSPSSLSASVGDRVTITCRASQSISSYLNWYQQKPGKAPKLLIYAASSLQSGVPSRFSGSGSGTDFTLTISSLQPEDFATYYCQQSYS--WTFGQGTKVEIK IGKV1-39*01, IGKJ1*01
DIQMTQSPSSLSASVGDRVTITCLASEGISSYLAWYQQKPGKAPKLLIYGANSLQTGVPSRFSGSGSATDYTLTISSLQPEDFATYYCQQSYKFPNTFGQGTKVEVK Humanized (Experimental)
DIQMTQSPASLSASLGETISIECLASEGISSYLAWYQQKPGKSPQLLIYGANSLQTGVPSRFSGSGSATQYSLKISSMQPEDEGDYFCQQSYKFPNTFGAGTKLELK Parental
DIQMTQSPSSLSASVGDRVTITCLASEGISSYLAWYQQKPGKAPKLLIYGANSLQTGVPSRFSGSGSGTDFTLTISSLQPEDFATYYCQQSYKFPNTFGQGTKLEIK Humanized (Sapiens^3)
DIQMTQSPSSLSASVGDRVTITCRASQSISSYLNWYQQKPGKAPKLLIYAASSLQSGVPSRFSGSGSGTDFTLTISSLQPEDFATYYCQQSYS--YTFGQGTKLEIK IGKV1-39*01, IGKJ2*01
```

#### Rovalpituzumab

##### Heavy chain

```
QVQLVQSGAEVKKPGASVKVSCKASGYTFTSYGISWVRQAPGQGLEWMGWISAYNGNTNYAQKLQGRVTMTTDTSTSTAYMELRSLRSDDTAVYYCAR-----YFDYWGQGTLVTVSS IGHV1-18*01, IGHJ4*01
QVQLVQSGAEVKKPGASVKVSCKASGYTFTNYGMNWVRQAPGQGLEWMGWINTYTGEPTYADDFKGRVTMTTDTSTSTAYMELRSLRSDDTAVYYCARIGDSSPSDYWGQGTLVTVSS Humanized (Experimental)
QIQLVQSGPELKKPGETVKISCKASGYTFTNYGMNWVKQAPGKGLKWMAWINTYTGEPTYADDFKGRFAFSLETSASTASLQINLKNEDTATYFCARIGDSSPSDYWGQGTTLTVSS Parental
QVQLVQSGSELKKPGASVKISCKASGYTFTNYGMNWVRQAPGQGLEWMGWINTYTGEPTYADDFKGRFVFSLDTSVSTAYLQISSLKAEDTAVYYCARIGDSSPSDYWGQGTLVTVSS Humanized (Sapiens^3)
QVQLVQSGSELKKPGASVKVSCKASGYTFTSYAMNWVRQAPGQGLEWMGWINTNTGNPTYAQGFTGRFVFSLDTSVSTAYLQISSLKAEDTAVYYCAR-----YFDYWGQGTLVTVSS IGHV7-4-1*02, IGHJ4*01
```

##### Light chain

```
EIVMTQSPATLSVSPGERATLSCRASQSVSSNLAWYQQKPGQAPRLLIYGASTRATGIPARFSGSGSGTEFTLTISSLQSEDFAVYYCQQYNN--WTFGQGTKVEIK IGKV3-15*01, IGKJ1*01
EIVMTQSPATLSVSPGERATLSCKASQSVSNDVVWYQQKPGQAPRLLIYYASNRYTGIPARFSGSGSGTEFTLTISSLQSEDFAVYYCQQDYTSPWTFGQGTKLEIK Humanized (Experimental)
SIVMTQTPKFLLVSAGDRVTITCKASQSVSNDVVWYQQKPGQSPKLLIYYASNRYTGVPDRFAGSGYGTDFSFTISTVQAEDLAVYFCQQDYTSPWTFGGGTKLEIR Parental
EIVMTQSPATLSVSPGDRVTITCKASQSVSNDVVWYQQKPGQAPKLLIYYASNRYTGVPSRFSGSGSGTDFTLTISSLQPEDFAVYYCQQDYTSPWTFGQGTKVEIK Humanized (Sapiens^3)
EIVMTQSPPTLSLSPGERVTLSCRASQSVSSYLSWYQQKPGQAPRLLIYGASTRATGIPARFSGSGSGTDFTLTISSLQPEDFAVYYCQQDYN--WTFGQGTKVEIK IGKV3-7*02, IGKJ1*01
```

#### Solanezumab

##### Heavy chain

```
EVQLLESGGGLVQPGGSLRLSCAASGFTFSSYAMSWVRQAPGKGLEWVSVIYSGGSSTYYADSVKGRFTISRDNSKNTLYLQMNSLRAEDTAVYYCAK-DYWGQGTLVTVSS IGHV3-23*03, IGHJ4*01
EVQLVESGGGLVQPGGSLRLSCAASGFTFSRYSMSWVRQAPGKGLELVAQINSVGNSTYYPDTVKGRFTISRDNAKNTLYLQMNSLRAEDTAVYYCASGDYWGQGTLVTVSS Humanized (Experimental)
EVKLVESGGGLVQPGGSLKLSCAVSGFTFSRYSMSWVRQTPEKRLELVAQINSVGNSTYYPDTVKGRFTISRDNAEYTLSLQMSGLRSDDTATYYCASGDYWGQGTTLTVSS Parental
EVQLVESGGGLVQPGGSLRLSCAASGFTFSRYSMSWVRQAPGKGLEWVAQINSVGNSTYYPDTVKGRFTISRDNAKNTLYLQMSSLRAEDTAVYYCASGDYWGQGTLVTVSS Humanized (Sapiens^3)
EVQLLESGGGLVQPGGSLRLSCAASGFTFSSYAMSWVRQAPGKGLEWVSVIYSGGSSTYYADSVKGRFTISRDNSKNTLYLQMNSLRAEDTAVYYCAK-DYWGQGTLVTVSS IGHV3-23*03, IGHJ4*01
```

##### Light chain

```
DVVMTQSPLSLPVTLGQPASISCRSSQSLVYSDGNTYLNWFQQRPGQSPRRLIYKVSNRDSGVPDRFSGSGSGTDFTLKISRVEAEDVGVYYCMQGTH--WTFGQGTKVEIK IGKV2-30*01, IGKJ1*01
DVVMTQSPLSLPVTLGQPASISCRSSQSLIYSDGNAYLHWFLQKPGQSPRLLIYKVSNRFSGVPDRFSGSGSGTDFTLKISRVEAEDVGVYYCSQSTHVPWTFGQGTKVEIK Humanized (Experimental)
DVVMTQTPLSLPVSLGDQASISCRSSQSLIYSDGNAYLHWFLQKPGQSPKLLIYKVSNRFSGVPDRFSGSGSGTDFTLKISRVETEDLGVYFCSQSTHVPWTFGGGTKLEIK Parental
DVVMTQTPLSLPVTLGQPASISCRSSQSLIYSDGNAYLHWYLQKPGQSPQLLIYKVSNRFSGVPDRFSGSGSGTDFTLKISRVEAEDVGVYYCSQSTHVPWTFGQGTKVEIK Humanized (Sapiens^3)
DVVMTQSPLSLPVTLGQPASISCRSSQSLVYSDGNTYLNWFQQRPGQSPRRLIYKVSNRDSGVPDRFSGSGSGTDFTLKISRVEAEDVGVYYCMQGTH--WTFGQGTKVEIK IGKV2-30*01, IGKJ1*01
```

### Talacotuzumab

#### Heavy chain

```
EVQLVQSGAEVKKPGESLRISCKGSGYSFTSYWISWVRQMPGKGLEWMGRIDPSDSYTNYSPSFQGQVTISADKSISTAYLQWSSLKASDTAMYYCAR-----AEYFQHWGQGTLVTVSS IGHV5-10-1*04, IGHJ1*01
EVQLVQSGAEVKKPGESLKISCKGSGYSFTDYYMKWARQMPGKGLEWMGDIIPSNGATFYNQKFKGQVTISADKSISTTYLQWSSLKASDTAMYYCARSHLLRASWFAYWGQGTMVTVSS Humanized (Experimental)
EVQLQQSGPELVKPGASVKMSCKASGYTFTDYYMKWVKQSHGKSLEWIGDIIPSNGATFYNQKFKGKATLTVDRSSSTAYMHLNSLTSEDSAVYYCTRSHLLRASWFAYWGQGTLVTVSA Parental
QVQLVQSGAEVKKPGASVKVSCKASGYTFTDYYMKWVRQAPGQGLEWMGDIIPSNGATFYNQKFKGRVTLTRDTSTSTAYMELNSLTSEDTAVYYCARSHLLRASWFAYWGQGTLVTVSS Humanized (Sapiens^3)
QVQLVQSGAEVKKPGASVKVSCKASGYTFTSYYMHWVRQAPGQGLEWMGIINPSGGSTSYAQKFQGRVTMTRDTSTSTVYMELSSLRSEDTAVYYCAR-----AEYFQHWGQGTLVTVSS IGHV1-46*01, IGHJ1*01
```

#### Light chain

```
DIVMTQSPDSLAVSLGERATINCKSSQSVLYSSNNKNYLAWYQQKPGQPPKLLIYWASTRESGVPDRFSGSGSGTDFTLTISSLQAEDVAVYYCQQYYS--YTFGQGTKLEIK IGKV4-1*01, IGKJ2*01
DIVMTQSPDSLAVSLGERATINCESSQSLLNSGNQKNYLTWYQQKPGQPPKPLIYWASTRESGVPDRFSGSGSGTDFTLTISSLQAEDVAVYYCQNDYSYPYTFGQGTKLEIK Humanized (Experimental)
DFVMTQSPSSLTVTAGEKVTMSCKSSQSLLNSGNQKNYLTWYLQKPGQPPKLLIYWASTRESGVPDRFTGSGSGTDFTLTISSVQAEDLAVYYCQNDYSYPYTFGGGTKLEIK Parental
DIVMTQSPDSLAVSLGERATINCKSSQSLLNSGNQKNYLTWYQQKPGQPPKLLIYWASTRESGVPDRFSGSGSGTDFTLTISSLQAEDVAVYYCQNDYSYPYTFGQGTKLEIK Humanized (Sapiens^3)
DIVMTQSPDSLAVSLGERATINCKSSQSVLYSSNNKNYLAWYQQKPGQPPKLLIYWASTRESGVPDRFSGSGSGTDFTLTISSLQAEDVAVYYCQQYYS--YTFGQGTKLEIK IGKV4-1*01, IGKJ2*01
```

### Tocilizumab

#### Heavy chain

```
QVQLQESGPGLVKPSETLSLTCTVSGGSI-SSHYWSWIRQPPGKGLEWIGYIYYSGSTNYNPSLKSRVTISVDTSKNQFSLKLSSVTAADTAVYYCAR------YFDYWGQGTLVTVSS IGHV4-59*11, IGHJ4*01
QVQLQESGPGLVRPSQTLSLTCTVSGYSITSDHAWSWVRQPPGRGLEWIGYISYSGITTYNPSLKSRVTMLRDTSKNQFSLRLSSVTAADTAVYYCARSLARTTAMDYWGQGSLVTVSS Humanized (Experimental)
DVQLQESGPVLVKPSQSLSLTCTVTGYSITSDHAWSWIRQFPGNKLEWMGYISYSGITTYNPSLKSRISITRDTSKNQFFLQLNSVTTGDTSTYYCARSLARTTAMDYWGQGTSVTVSS Parental
QVQLQESGPGLVKPSQTLSLTCTVSGYSITSDHAWSWIRQPPGKGLEWIGYISYSGITTYNPSLKSRVSISVDTSKNQFSLKLNSVTAADTAVYYCARSLARTTAMDYWGQGTLVTVSS Humanized (Sapiens^3)
QVQLQESGPGLVKPSETLSLTCTVSGGSI-SSHYWSWIRQPPGKGLEWIGYIYYSGSTNYNPSLKSRVTISVDTSKNQFSLKLSSVTAADTAVYYCAR------YFDYWGQGTLVTVSS IGHV4-59*11, IGHJ4*01
```

#### Light chain

```
DIQMTQSPSSLSASVGDRVTITCQASQDISNYLNWYQQKPGKAPKLLIYDASNLETGVPSRFSGSGSGTDFTFTISSLQPEDIATYYCQQYDN--WTFGQGTKVEIK IGKV1-33*01, IGKJ1*01
DIQMTQSPSSLSASVGDRVTITCRASQDISSYLNWYQQKPGKAPKLLIYYTSRLHSGVPSRFSGSGSGTDFTFTISSLQPEDIATYYCQQGNTLPYTFGQGTKVEIK Humanized (Experimental)
DIQMTQTTSSLSASLGDRVTISCRASQDISSYLNWYQQKPDGTIKLLIYYTSRLHSGVPSRFSGSGSGTDYSLTINNLEQEDIATYFCQQGNTLPYTFGGGTKLEIN Parental
DIQMTQSPSSLSASVGDRVTITCRASQDISSYLNWYQQKPGKAPKLLIYYTSRLHSGVPSRFSGSGSGTDFTLTISSLQPEDFATYYCQQGNTLPYTFGQGTKLEIK Humanized (Sapiens^3)
DIQMTQSPSSLSASVGDRVTITCRASQSISSYLNWYQQKPGKAPKLLIYAASSLQSGVPSRFSGSGSGTDFTLTISSLQPEDFATYYCQQSYS--YTFGQGTKLEIK IGKV1-39*01, IGKJ2*01
```
